# The efficacy and safety of *Momordica charantia* L. in animal models of type 2 diabetes mellitus; A systematic review and meta-analysis

**DOI:** 10.1101/681494

**Authors:** Emanuel L. Peter, Prakash B. Nagendrappa, Anita Kaligirwa, Patrick Engeu Ogwang, Crispin Duncan Sesaazi

## Abstract

**Background:** *Momordica charantia* L. (Cucurbitaceae) has been used to control hyperglycemia in people with type 2 diabetes mellitus in Asia, South America, and Africa for decades. However, a meta-analysis of clinical trials confirmed very low-quality evidence of its efficacy. To potentially increase the certainty of evidence, we evaluated the effect of *M. charantia* L. in comparison with vehicle on glycemic control in animal models of type 2 diabetes mellitus.

**Methods:** Review authors searched in MEDLINE, Web of Science, Scopus, and CINAHL databases without language restriction through April 2019. Two authors independently evaluated full texts, assessed the risk of bias, and extracted data. We analyzed the influence of study design and evidence of publication bias.

**Results:** The review included 66 studies involving 1861 animals. They had a follow up between 7 and 90 days. Majority 29 (43.9%) used Wistar albino rats, and 37 (56.1%) used male animals. Thirty-two (48%) used an aqueous extract of fresh fruits. *M. charantia* L. reduced fasting plasma glucose (FPG) and glycosylated hemoglobin A1c in comparison to vehicle control (42 studies, 815 animals; SMD, −6.86 [95% CI; −7.95, −5.77], 3 studies, 59 animals; SMD; −7.76 [95%CI; −12.50, −3.01]) respectively. Magnitude of FPG was large in Wistar albino rat subgroup; SMD; −10.29, [95%CI; −12.55, −8.03]. Publication bias changed FPG to non-significant −2.46 SMD, [95%CI; - 5.10, 0.17]. We downgraded the evidence to moderate quality due to poor methodological quality, high risk of bias, unexplained heterogeneity, suspected publication bias, and lack of standardized dose.

**Conclusion:** *M. charantia* L. lowers elevated plasma glucose level in type 2 diabetes mellitus animal models. Publication bias and poor methodological quality call for future researches to focus on standardizing dose with chemical markers and provide measures to improve preclinical type 2 diabetes mellitus studies.

Systematic review registration CRD42019119181

## Introduction

Type 2 diabetes mellitus (T2DM) is a chronic hyperglycemic condition in response to progressively impaired glucose regulation due to insulin resistance and beta-cell dysfunction [1, 2]. The chronicity of hyperglycemia causes microvascular complications in the retina, renal glomerulus and peripheral nerves [3]; and increases the risk of accelerated atherosclerosis and premature death [4]. Other complications include dementia, sexual dysfunction, depression and lower-limb amputations [5–7].

People with T2DM use oral hypoglycemic agents (OHAs) for glycemic control and ameliorating diabetes complications, but the OHAs have in recent years been linked to intolerable side effects and increasing failure rate [8], leaving the majority of people with T2DM using medicinal plants as alternative therapy [9, 10]. *Momordica charantia* L. (Family; Cucurbitaceae) is one of such medicinal plants and well known in African, Ayurveda, and traditional Chinese systems of medicine for its use in diabetes mellitus. It is also a vital market vegetable in southern and eastern Asia, and most African countries [11, 12].

The antidiabetic activity of *M. charantia* L. has been investigated in several animal models of type 2 diabetes mellitus [13, 14]. Majority of these studies used chemically induced T2DM in various animal species and assessed improvement of features of T2DM such as hyperglycemia, insulin resistance, beta-cell dysfunction, serum insulin level, beta-cell mass, dyslipidemia [15–18]. These features are also crucial in the clinical evaluation of the efficacy of antidiabetic activity of herbal products as reported in previous systematic reviews and meta-analysis based on randomized clinical trials [19, 20].

Pharmacological studies have established several potential modes of actions through which *M. charantia* L. could lower high blood glucose and prevent complications. The proposed mechanisms include; improved histological architecture of the islets of Langerhans and beta-cell regeneration [16,21,22], insulin secretagogue [23, 24], enhance peripheral glucose utilization, inhibit glucose-6-phosphatase and fructose biphosphatase glucogenic enzymes [25], increases peroxisome proliferator-activated receptor gamma (PPAR-γ) activity and decreases protein Kinase C (PKC-β) activity in kidneys [26]. Despite the number of preclinical studies performed each year continuing to increase, and our understanding of *M. charantia* L. mode of action is improving, a recent meta-analysis of five randomized clinical trials confirmed its glucose lowering ability with only very low certainty of evidence [19]. There is also a lack of consensus on the proposed mode of action due to contradictory findings of existing preclinical studies. The contradictory findings of preclinical studies and the weak clinical evidence indicate the existing challenges in translating animal studies of *M. charantia* L. to clinical practice. Given the sheer volume of preclinical experiments of the efficacy of treatment with *M. charantia* L. in T2DM, a structured process is needed to objectively evaluate and provide robust, informative summaries of these studies.

Systematic review and meta-analysis of animal studies is one of the promising structured process of assessment which could facilitate rigorous methodological quality, risk of bias, and publication bias assessment in animal studies and determine their influence on clinical generalizability of animal studies findings [27–29]. Given the very low-quality human evidence, considering evidence from animal studies might change the assessment of the apparent magnitude of effect or might potentially increase our certainty in the evidence [30].

Therefore, this systematic review and meta-analysis of animal studies aimed to evaluate the evidence of the efficacy of treatment with *M. charantia* L. on animal models of type 2 diabetes mellitus. We also described the impact of study design methodological features and publication bias on the efficacy of *M. charantia* L. and suggested areas of focus for future animal research which may reliably predict clinical efficacy.

## Materials and methods

This systematic review and meta-analysis is based on registered protocol number CRD42019119181 [31]. We reported results according to the PRISMA guidelines, the PRISMA abstract checklist, and guidelines for reporting systematic review and meta-analysis of animals studies [28,32,33].

### Information source and search strategy

Review authors searched MEDLINE through PubMed platform, Web of Science through a web of knowledge platform, CINAHL, and Scopus. The authors also searched gray literature to include conference papers, technical reports, thesis and Dissertations in Google Scholar, Google, OpenGrey, ProQuest Dissertations & Theses, and British Library EThos. Review authors searched each database through April 2019. They also screened reference lists of included studies and reviews for additional eligible studies not retrieved by the search.

The search strategy involved a combination of MeSH terms and keywords. The search terms were divided into three components i.e., the population component with the following words; “animals,” “animal,” “animals model,” “preclinical studies,” “experimental animals,” “experimental animal,” “laboratory animal,” “laboratory animals,” “rodents,” “rodent,” “rabbits,” “rabbit,” “rats,” “rat,” “diabetic rats,” “animal disease model,” “mice,” “mouse.” The intervention component’s terms were “*Momordica charantia,” “*bitter melon,” “bitter gourd,” and “karela.” The last component had “diabetes mellitus, type 2,” “non-insulin dependent diabetes mellitus,” “NIDDM,” “glucose metabolic disorders,” “metabolic diseases,” “hyperlipidemia,” “hyperglycemia,” “insulin resistance,” and “glucose intolerance” terms. The three search components were combined with the boolean logic term “AND” while the keywords within each component were combined with “OR.” Search filters for the identification of preclinical studies in PubMed were applied to increase search efficiency [34]. Review authors did not restrict language during the search and identification of studies. The final searches for each database were re-run just before the final analyses to retrieve the most recent studies eligible for inclusion: The appendix S2 elaborated search strategy and their results for PubMed, Scopus, and CINAHL databases (S2 Appendix).

### Study design and animal models eligibility

Review authors included experimental animal studies if they were either randomized or non-randomized controlled designed, original full article with data presented numerically or graphically, and those conducted in animal models of type 2 diabetes mellitus. The animal models were carefully assessed to include those which closely mimic at least some aspects of the pathophysiology of humans with type 2 diabetes mellitus such as insulin resistance and β-cells failure to ensure construct validity [35]. Our review also included all sex, age, species and strain of animals. However, the review excluded studies done in a human, *in vitro*, *ex vivo,* and *in-silico* designs, and before-after studies without a description of the control group.

### Intervention and comparison eligibility

The preclinical intervention group included animals from studies that evaluated the efficacy or safety of the treatment with *M. charantia* L. preparations (whole extract or fraction of any part of the *M. charantia* L.) in any dosing, dosage forms, and frequency. The included studies should have induced T2DM in animals before administered the *M. charantia* L. preparations. The comparison group included animals from studies that induced experimental T2DM and treated with vehicle or standard treatment. Healthy animal control was also included to establish the extent of T2DM induction. The review excluded preclinical studies that evaluated the efficacy of polyherbal preparations of *M. charantia* L. or isolated pure compounds, concurrent treatment with standard oral hypoglycemic agents or insulin, and control treated with any other drug.

### Study records

#### 1. Study selection and data management

Review authors pooled identified articles into Mendeley software var. 2.1 (Elsevier). After deduplication, the titles and abstracts of studies retrieved using the search strategy and those from additional sources were screened independently by two review authors (ELP & AK). The two authors then retrieved full texts and independently assessed them for eligibility against predetermined inclusion criteria. They resolved their disagreement over the eligibility of particular studies through consensus.

#### 2. Data items and collection process

Two review authors extracted data independently from the included studies using a pilot tested data collection form. Discrepancies between the authors were identified and resolved through consensus. Reviewers contacted corresponding authors of included studies via email to obtain numerical data of studies that had data presented graphically, missing or when additional data were required. Two categories of data of interest extracted were; 1) Primary outcome representing fasting plasma glucose level, and 2) Secondary outcomes included; glycosylated hemoglobin A1c (HBA1c), Homeostatic Model Assessment of Insulin Resistance (HOMA-IR), Homeostatic model assessment for assessing β-cell function (HOMA-B), serum insulin level, number of insulin-positive cells, triglycerides (TGs), total cholesterol (TC), high density lipoprotein cholesterol (HDL-c), low density lipoprotein cholesterol (LDL-c), liver glycogen, weight, alanine aminotransferase (ALT), aspartate aminotransferase (AST), alkaline phosphatase (ALP), gamma-glutamyl transpeptidase (GGT), urea, and serum creatinine.

### Taxonomical assessment

The taxonomical and nomenclatural accuracy was assessed by comparing reported taxonomical information with existing standards in open botanical database accessible at www.theplantlist.org. Frequency of erroneous names use, types of such errors, identification of a specimen, and voucher specimen deposited were assessed according to methods proposed by Rivera and colleagues [36]. The authors gave “A” grade for studies with full information about the species of plant, identification of specimen, and deposited voucher specimen, while they grade “B” those studies with partial information about the species of plant such as studies which did not present information on identification of specimen and a voucher specimen and those with inaccurate taxonomic information. Finally, the authors rated “C” to studies with incomplete or not presented at all information about the species of plant, or identification of specimens and a voucher specimen.

### Methodological quality and risk of bias assessment

Review authors used SYRCLE’s risk of bias tool to assess the risk of bias for each preclinical animal study included [37]. The tool assessed domains of random sequence generation, baseline characteristics, allocation concealment, random housing, blinding of investigators/caregivers, random outcome assessment, blinding of assessor, incomplete outcome data, selective outcome reporting, and other sources of bias. Each criterion was assigned value as high, low or unclear risk of bias. The authors also used a modified CAMARADES checklist to assess the methodological quality of the included studies. This checklist combined the reporting of several measures to reduce bias and some indicators of external validity. The quality indicators are based on 10 criteria; 1) peer-reviewed publication 2) statement of control of temperature 3) random allocation to treatment or control 4) blinded caregiver/investigator 5) blinded assessment of outcome 6) use of co-interventions/co-morbid 7) appropriate animal model (age, sex, species, strain) 8) sample size calculation 9) compliance with animal welfare regulations 10) statement of potential conflict of interests [38]. Each study was given a quality score out of a possible total of 10 points. Finally, the authors calculated mean score and categorized studies into “low quality” for mean score 1–5 and “high quality” for mean score 6–10.

### Data synthesis

Quantitative data were pooled in a statistical meta-analysis using Review Manager (RevMan) software 5.3 (Copenhagen: The Nordic Cochrane Centre, The Cochrane Collaboration, 2014). Continuous variables analyzed in a meta-analysis included; FPG, HbA1c, serum insulin level, number of insulin-positive cells, TGs, TC, HDL-c, LDL-c, liver glycogen, ALT, AST, ALP, urea, serum creatinine, and weight. Since the same outcomes reported in different measurement scale, we used the standardized mean difference (SMD) to evaluate the effect of *M. charantia* L. in comparison to vehicle control. The SMD considered the difference in means between intervention and control groups at follow up divided by pooled standard deviation of the two groups to convert all outcome measures to a standardized scale with a unit of standard deviation. The inverse of variance-weighted method was used to attribute the relative contribution of each included study to pooled SMD effect of *M. charantia* L. and its 95% confidence intervals [39]. The authors used the random effect model for pooling effect estimates because the effect sizes from animal studies were more likely to differ due to the difference in design characteristics.

Qualitative data were summarized in the form of a table. We used signs (+) and (-) to indicate the direction of increased or decreased effect respectively. Variables analyzed qualitatively were HOMA-IR, HOMA-B, morphological structure of islet of Langerhans, number of beta-cells and number of insulin secretory granules.

#### 1. Heterogeneity assessment

We used the I^2^ statistic to quantify heterogeneity in primary studies [40]. The I^2^ of 75 or more was considered as indicative of substantial heterogeneity [41, 42]. Sensitivity analysis was done to examine potential factors that influence heterogeneity on the primary outcome (FPG). For this analysis we considered risk of bias score, methodological quality score, and performed subgroups analysis by study design (randomized and non-randomized design), duration of treatment, dose, mode of preparation of *M. charantia* L., animal species (mouse, rat, rabbit, dog, other), animal strains (KK mice, C57BL/6J mice, others), animal age, sex (male, female), and model of induction of type 2 diabetes mellitus (chemical, genetic, surgical, high-fat diet).

#### 2. Publication bias

Publication bias for each outcome was assessed by testing the asymmetry of the funnel plot using Egger’s test [43]. For the publication bias assessment, we only considered meta-analysis of ten or more studies because test power is generally too low to distinguish chance from real asymmetry when it includes a smaller number of the primary studies [43, 44]. When publication bias was detected, the trim and fill method was used to correct the probable publication bias by imputing missed studies and adjusted the effect size [45].

#### 3. Assessment of confidence in cumulative evidence

Review authors used “The Grading of Recommendations, Assessment, Development, and Evaluation (GRADE) approach” as a framework to rate the certainty in the evidence of preclinical animal studies [30, 46]. The authors rated the certainty for each outcome by considering the risk of bias (as assessed by SYRCLE’S risk of bias tool), inconsistency (as assessed by heterogeneity tests, confidence intervals, and P-values), imprecision, publication bias, and indirectness as proposed by Leeflang and colleagues [30]. After considering all factors, the authors rated evidence as high, moderate, low or very low-quality.

## Results

### Results of the search

We identified 443 articles through electronic and manual searching. After removing duplications and screening the articles based on the titles and abstracts, 181 articles remained. The full-texts of these articles were examined for eligibility according to the inclusion and exclusion criteria. Review authors further excluded 115 articles because one was a thesis for which a published article retrieved, 12 were only abstracts, 16 were inappropriately designed, 45 had not induced T2DM before administering the *M. charantia* L., 18 had no outcomes of interest, 16 did not investigate the intervention of interest and seven were duplicate publications. For each set of duplicate publications, we included one article which had most data. We also contacted the corresponding authors, and only one responded and shared full texts. The remaining majority of other authors failed to respond. Therefore, the systematic review qualitatively synthesized graphically presented data. Finally, we included 66 studies in qualitative analysis and 48 studies in meta-analysis. A PRISMA flow diagram is presented to show the screened, excluded and included articles (Fig 1).

**Fig 1.**
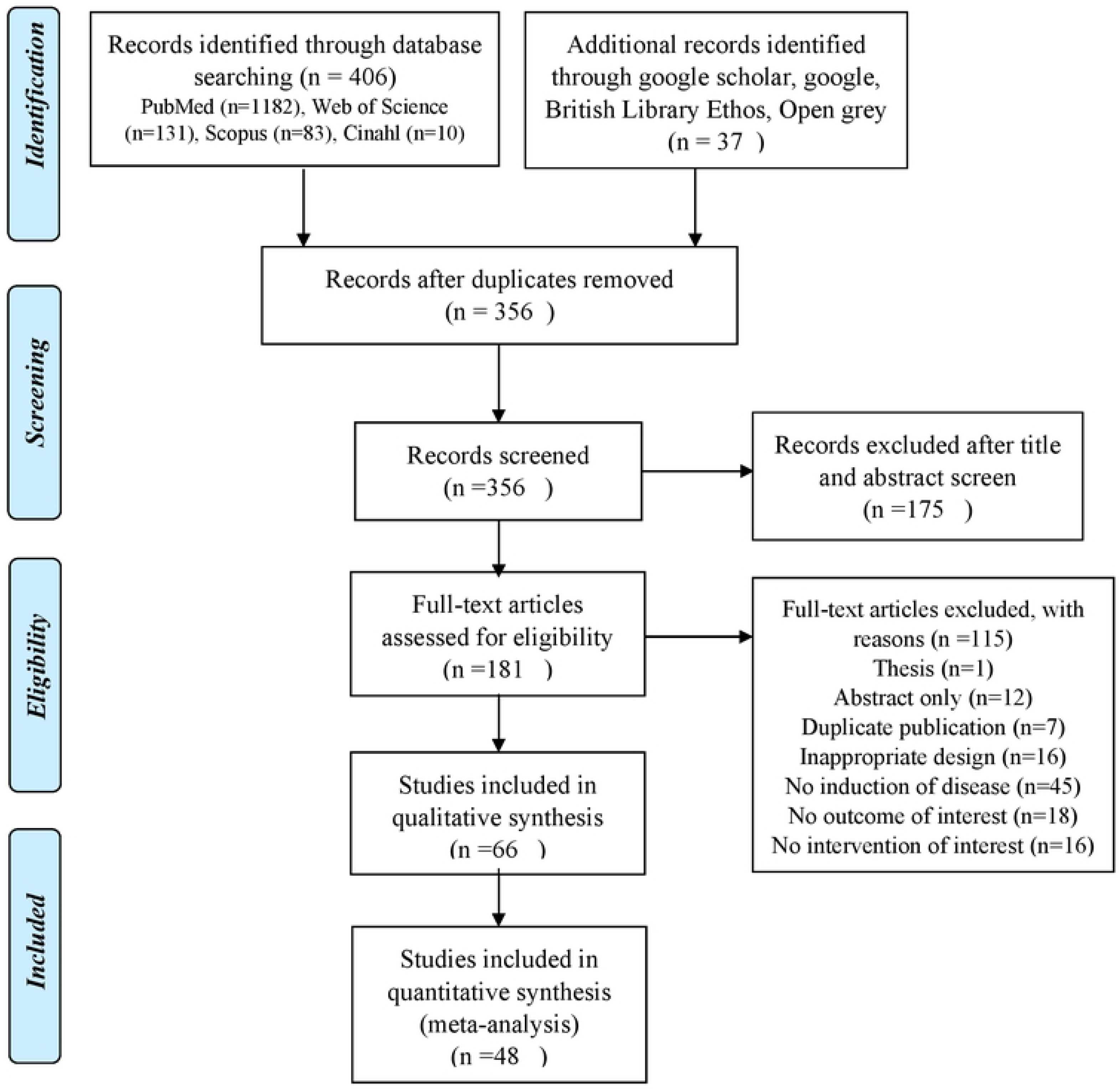
Flow diagram for screened included and excluded studies.

### Description of included studies

The majority of included studies 51 (77.3%) used rats, whereas 12 (18.2%) used mice and only 3 (4.5%) used rabbits. Regarding the strains of animal species used; 29 (43.9%) studies used Wistar albino rats, 11 (16.7%) used Sprague-Dawley rats and the remaining 14 different strains used are as shown in table 1. However, three of the included studies did not specify any strains used.

**Table 1.**
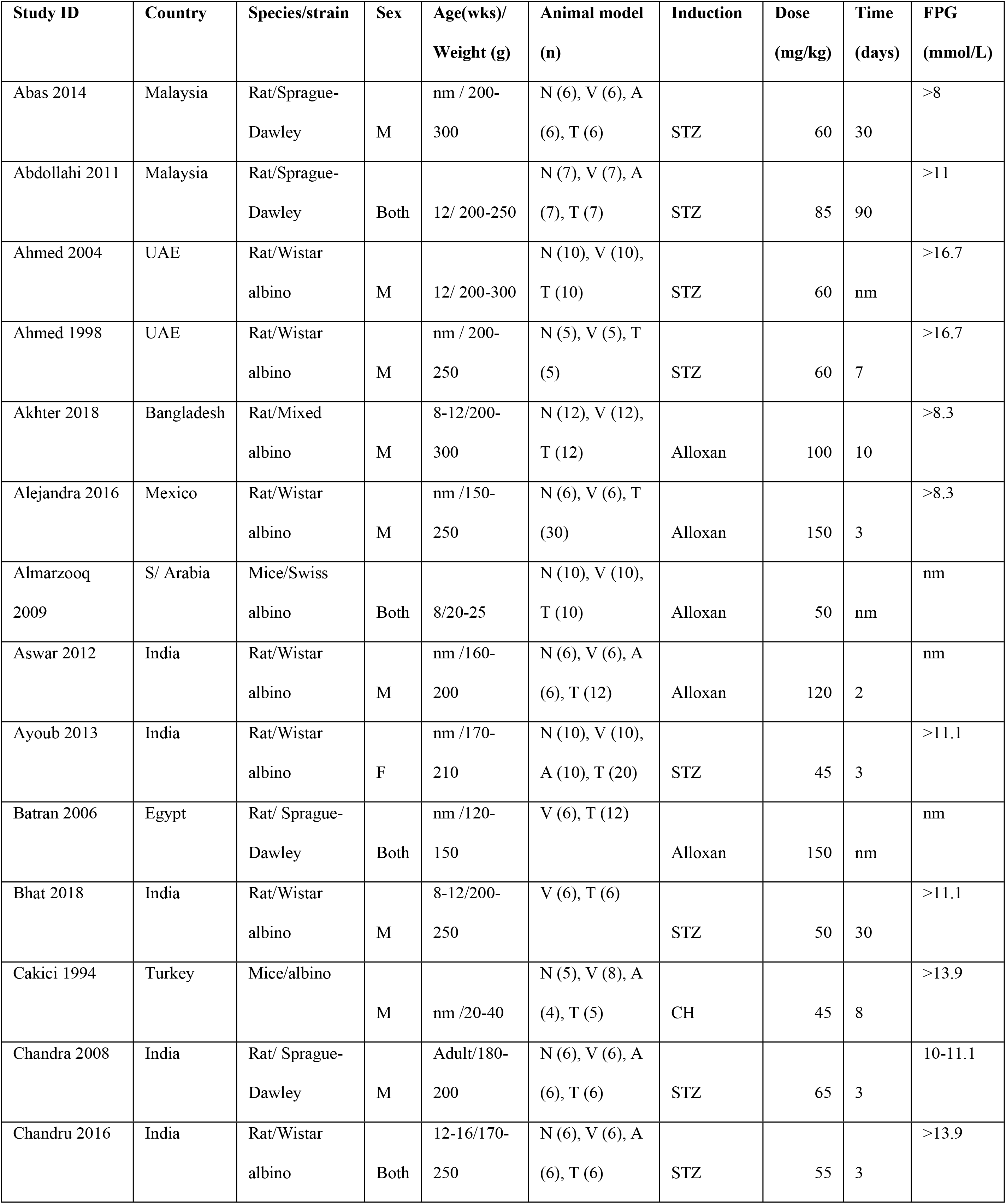
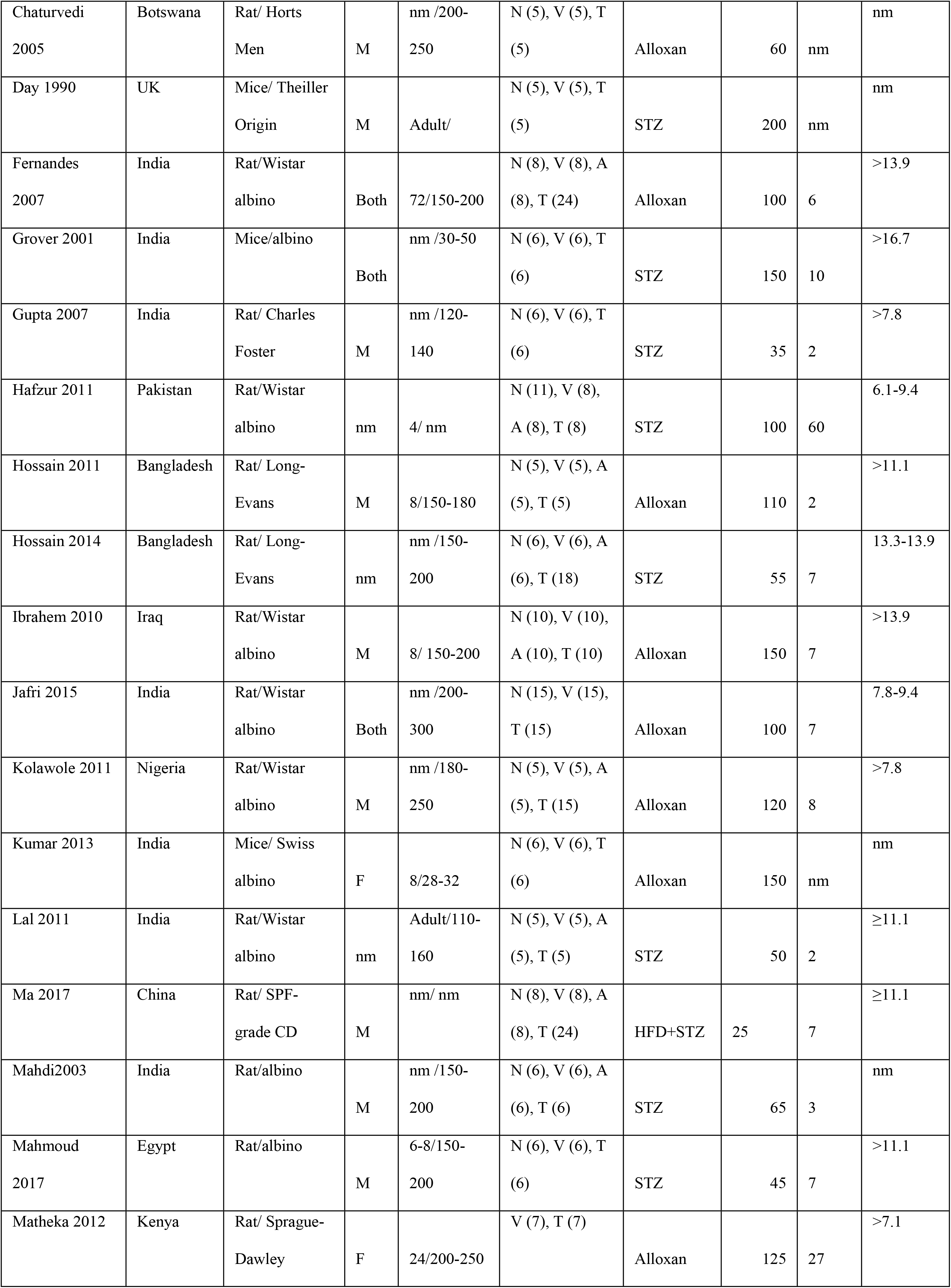
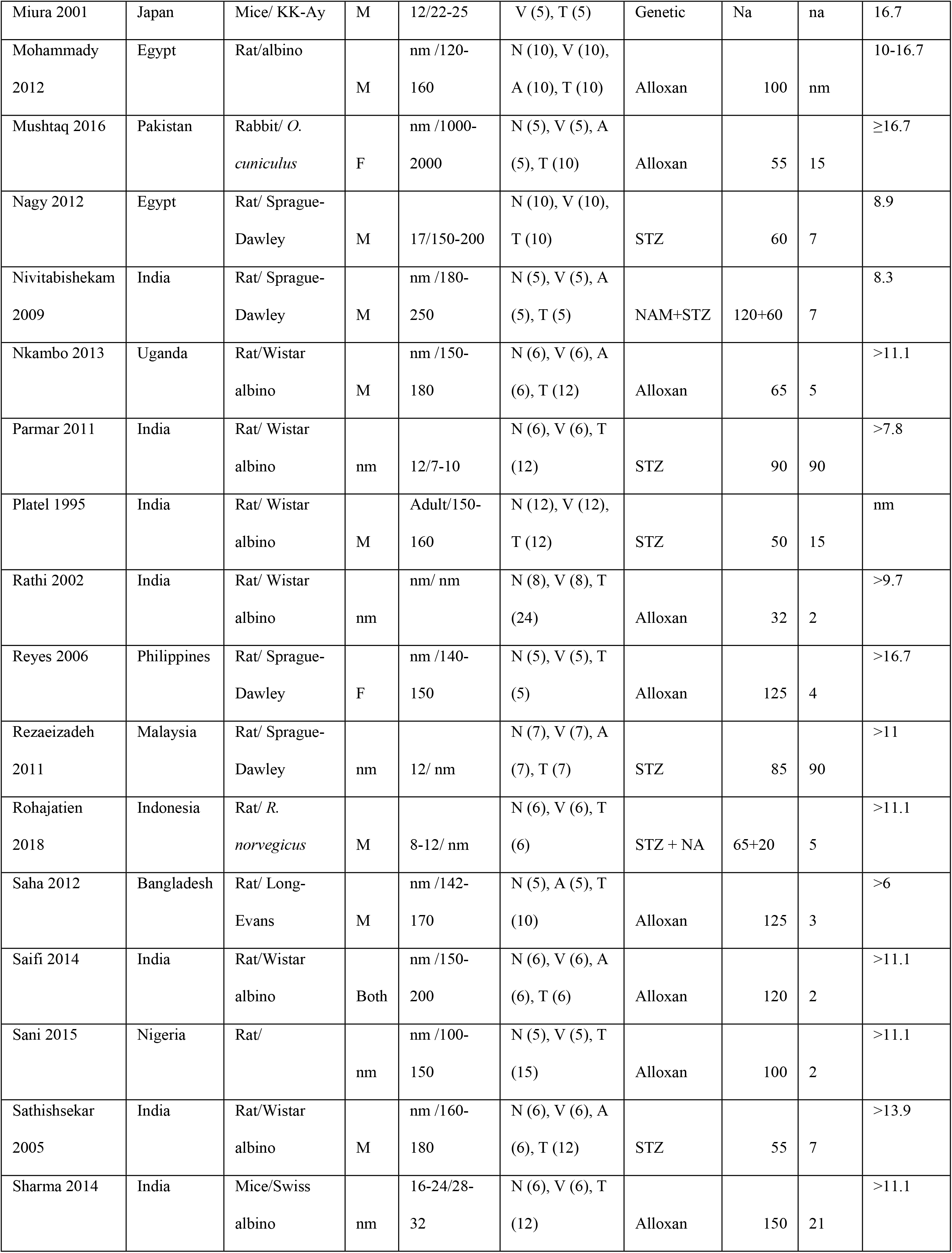
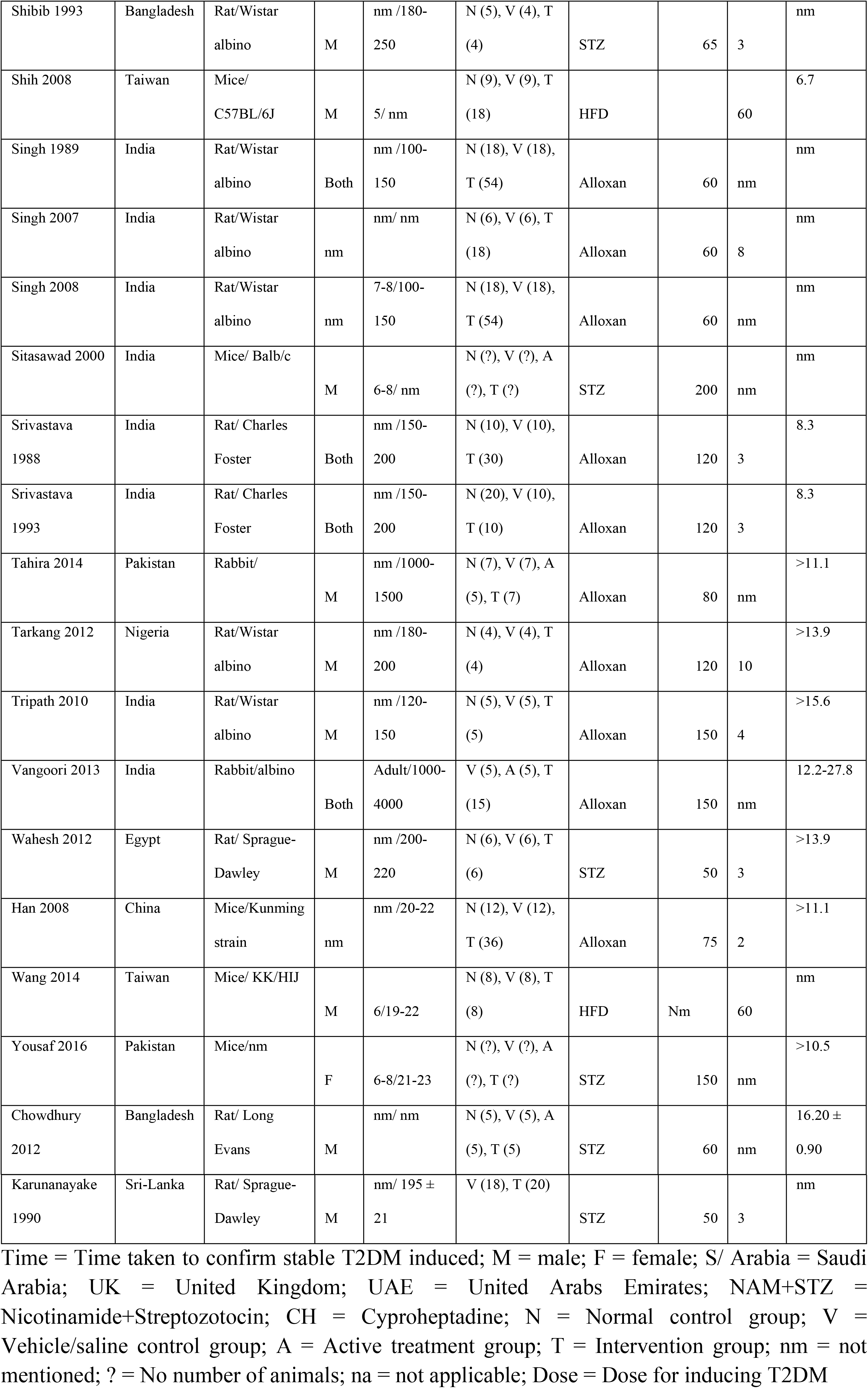
Baseline characteristics of the included studies.

Among the studies included, 37 (56.1%) used male animals, six used only female animals (9.1%), 12 (18.2%) used equal numbers of male and female animals, and the remaining studies did not provide information on the sex of the animals used.

About 32 (48.5%) studies used alloxan monohydrate, 27 (40.9%) studies used Streptozotocin (STZ) whereas two studies each used a high-fat diet and nicotinamide + STZ. One study each used remaining induction materials; cyproheptadine, genetically induced model, and high-fat diet + STZ.

The dose of STZ used ranges from a minimum of 35 mg/kg to a maximum of 200 mg/kg. The average dose was 77.59 ± 44.9 mg/kg. On the other hand, alloxan monohydrate used at a minimum dose of 32 and a maximum dose of 150 mg/kg with an average of 104.75 ± 35.447 mg/kg. The time taken to confirm stable T2DM after exposure to induction material was between one and 90 days. The majority of studies used models of T2DM with FPG levels at baseline ≥ 11.1 mmol/L, while three studies had models with ≥ 6 mmol/L [47–49].

Thirty-two (32) of these studies used an aqueous extract of fresh or dried fruits, and 17 used an alcoholic extract. The remaining studies used acetone extract, hydroalcoholic extract, petroleum ether extract, supernatant aqueous extract, and powdered dried fruits.

About 62 studies used fruits of *M. charantia* L., three used leaves, and one used seeds. Saifi et al., 2014 is the only study that described quality control measures of the intervention; the remaining studies did not describe quality control measures (Table 2). The studies administered the *M. charantia* L. between 7 days and 90 days. Table 1 & 2 summarized characteristics of the included studies.

**Table 2.**
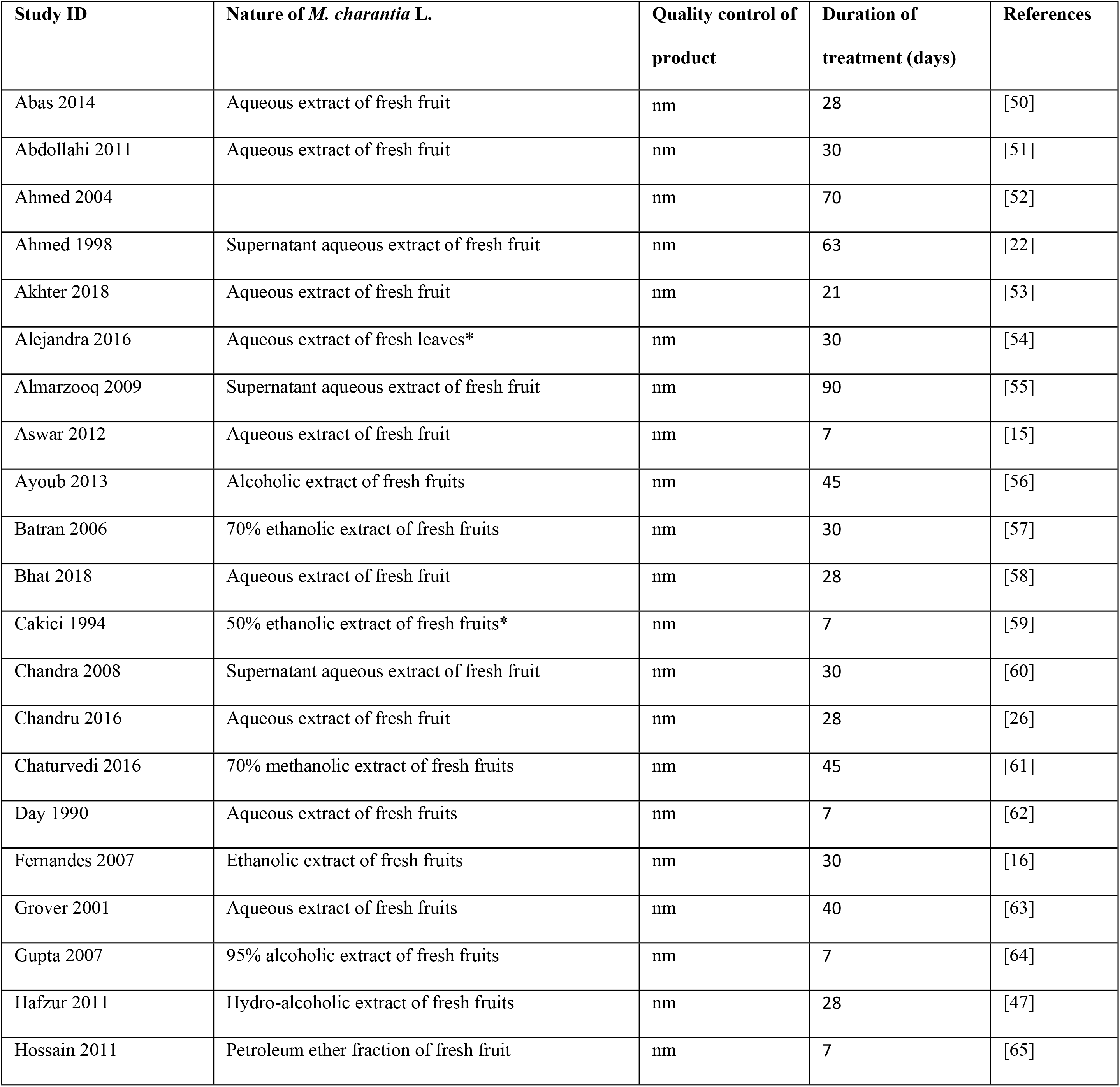
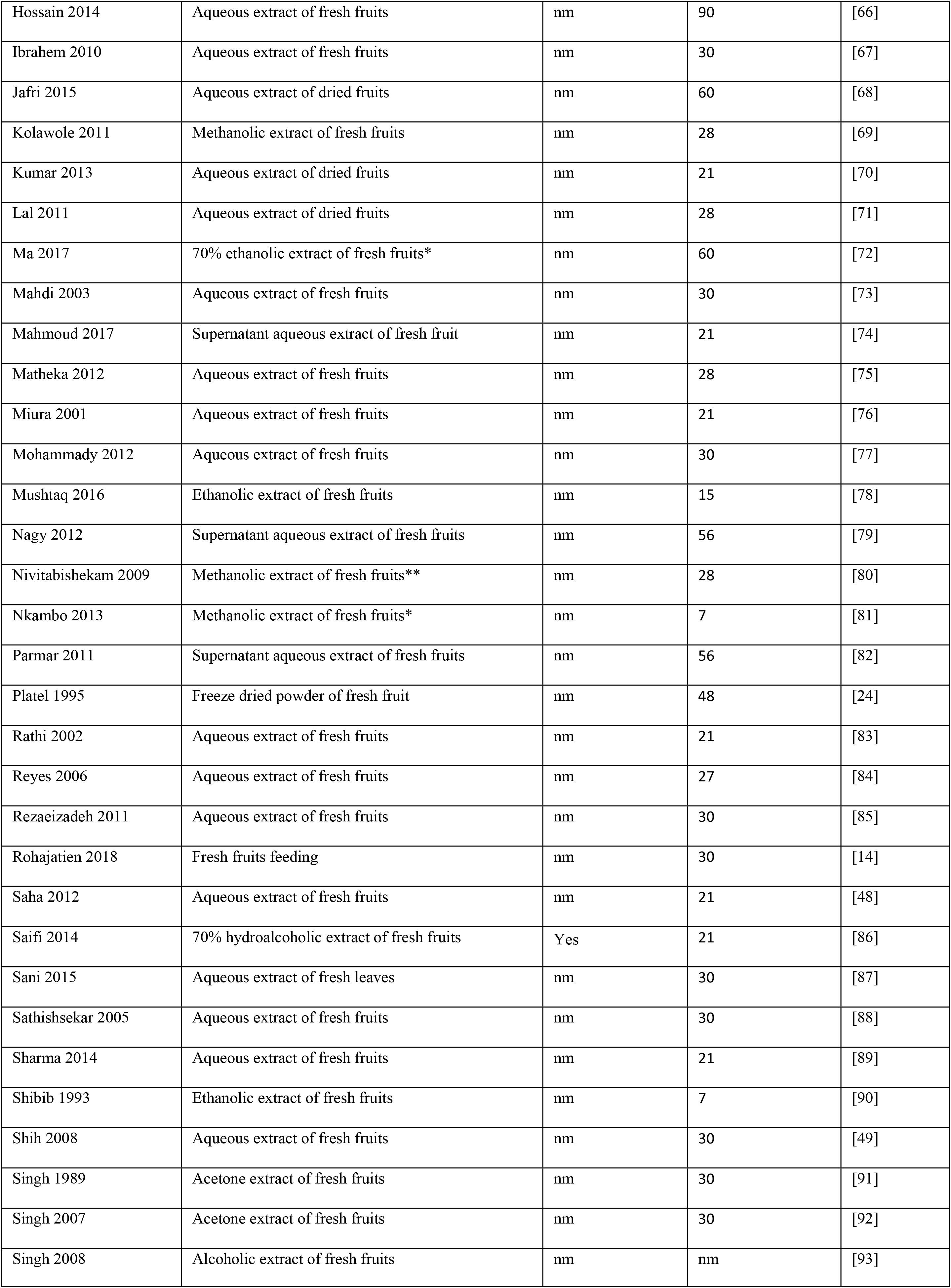
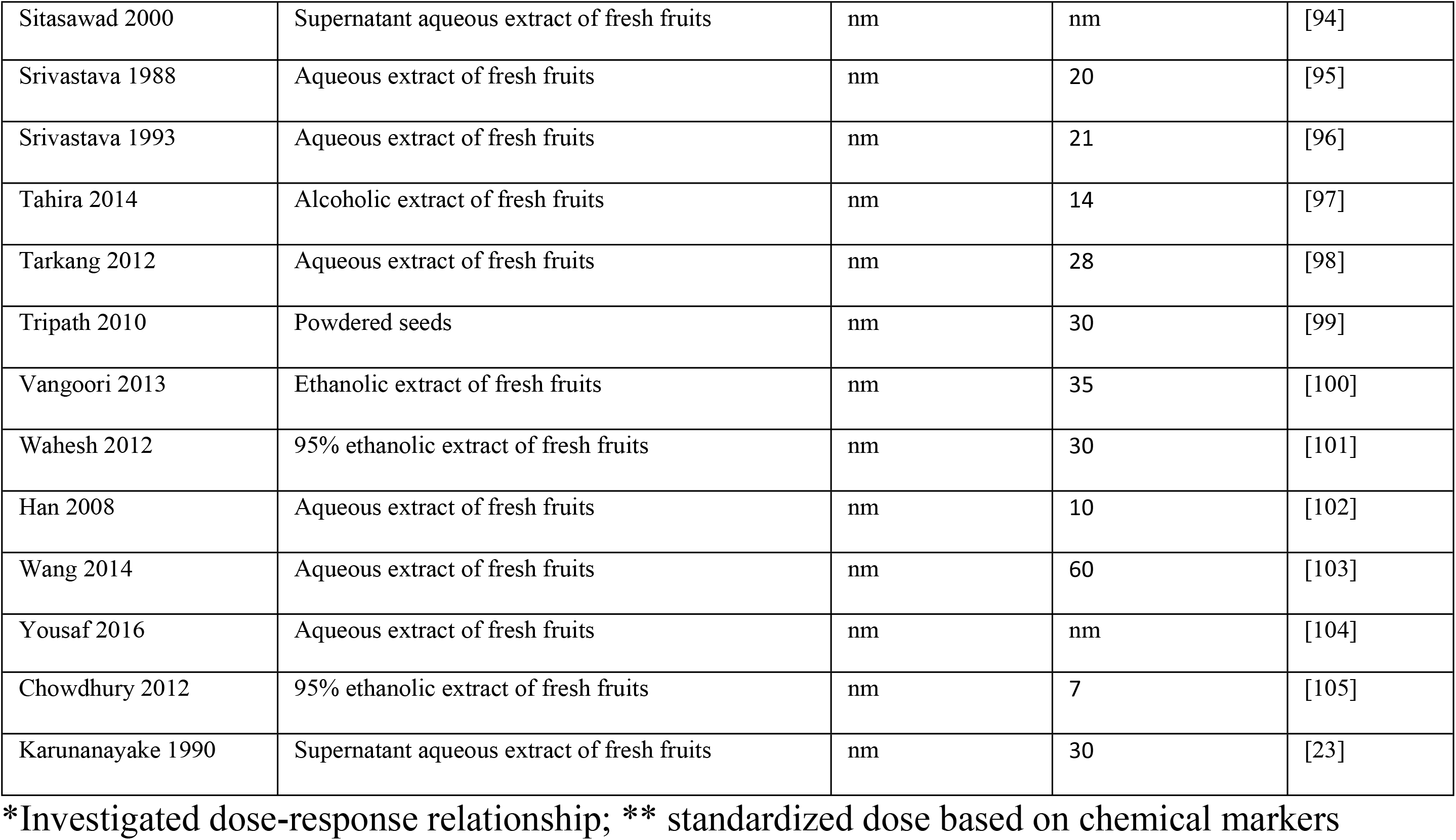
Baseline characteristics of the included studies.

### Taxonomical assessment of included studies

All 66 included studies used the scientific names; however, the majority 58 (87.9%) of the scientific names were not correct. The most recurrent type of error was missing plant authority names 39 (59.9%) and missing plant family names 26 (39.4%). Table 3 illustrates other types of errors identified.

**Table 3.** The common type of errors in scientific name identified in the included studies.

| Particulars | N (%) |
| --- | --- |
| Use of the scientific name |  |
| Correct name | 8 (11.6) |
| Incorrect name | 58 (87.9) |
| Type of errors in incorrect names |  |
| Misspelling of genus or species | 3 (5.2) |
| Inconsistent use of Italics | 6 (10.3) |
| Inappropriate capitalization | 5 (8.6) |
| Missing author | 39 (59.9) |
| Non-standard author abbreviation | 13 (22.4) |
| Missing of a period of abbreviation | 5 (8.6) |
| No family name | 26 (39.4) |
| Misspelling | 7 (12.1) |

Four (4) out of 66 included studies were given taxonomical validation score of “A” because they presented full information about plant name, identification of specimens, and voucher specimen deposited. On the other hand, ten studies were given a score of “B” since only partial information about plant name and identification of specimen were present. It is worth noting that, the majority of included studies (52) had inadequate or no information about taxonomical identification of plant species (Table 4).

**Table 4.**
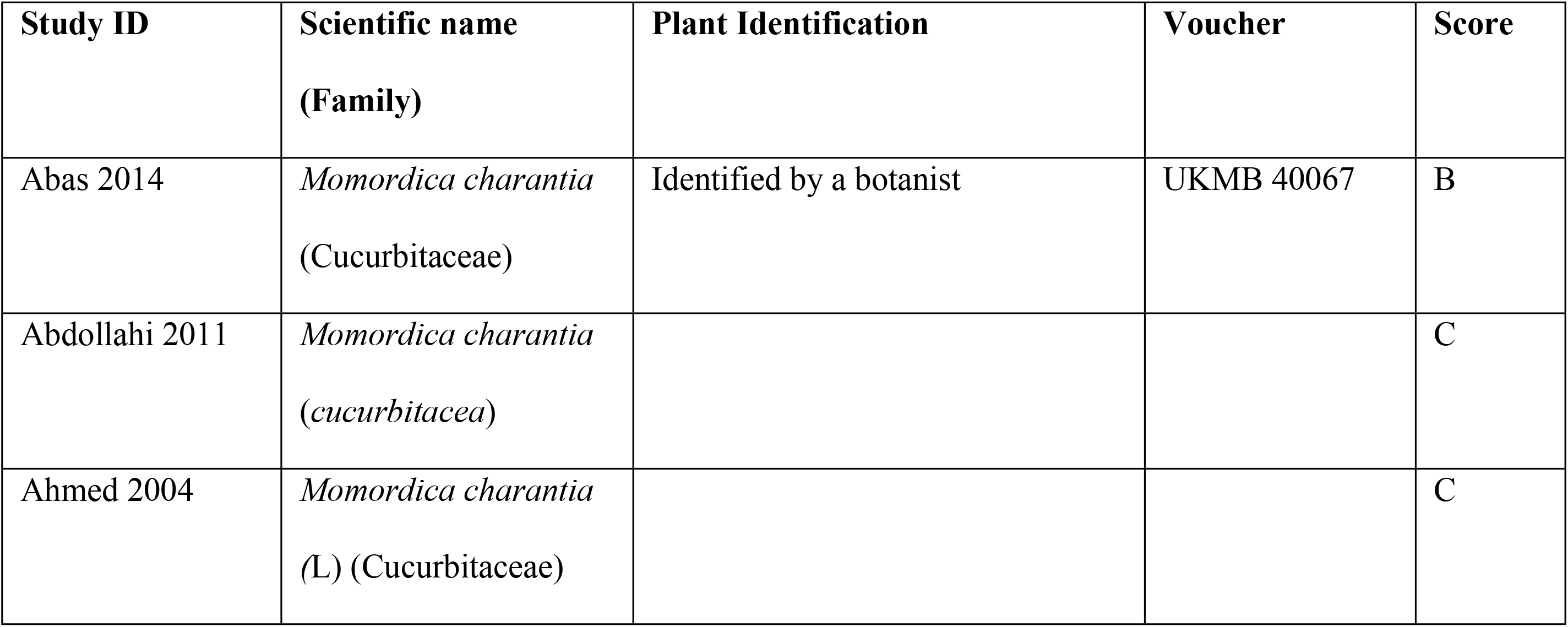
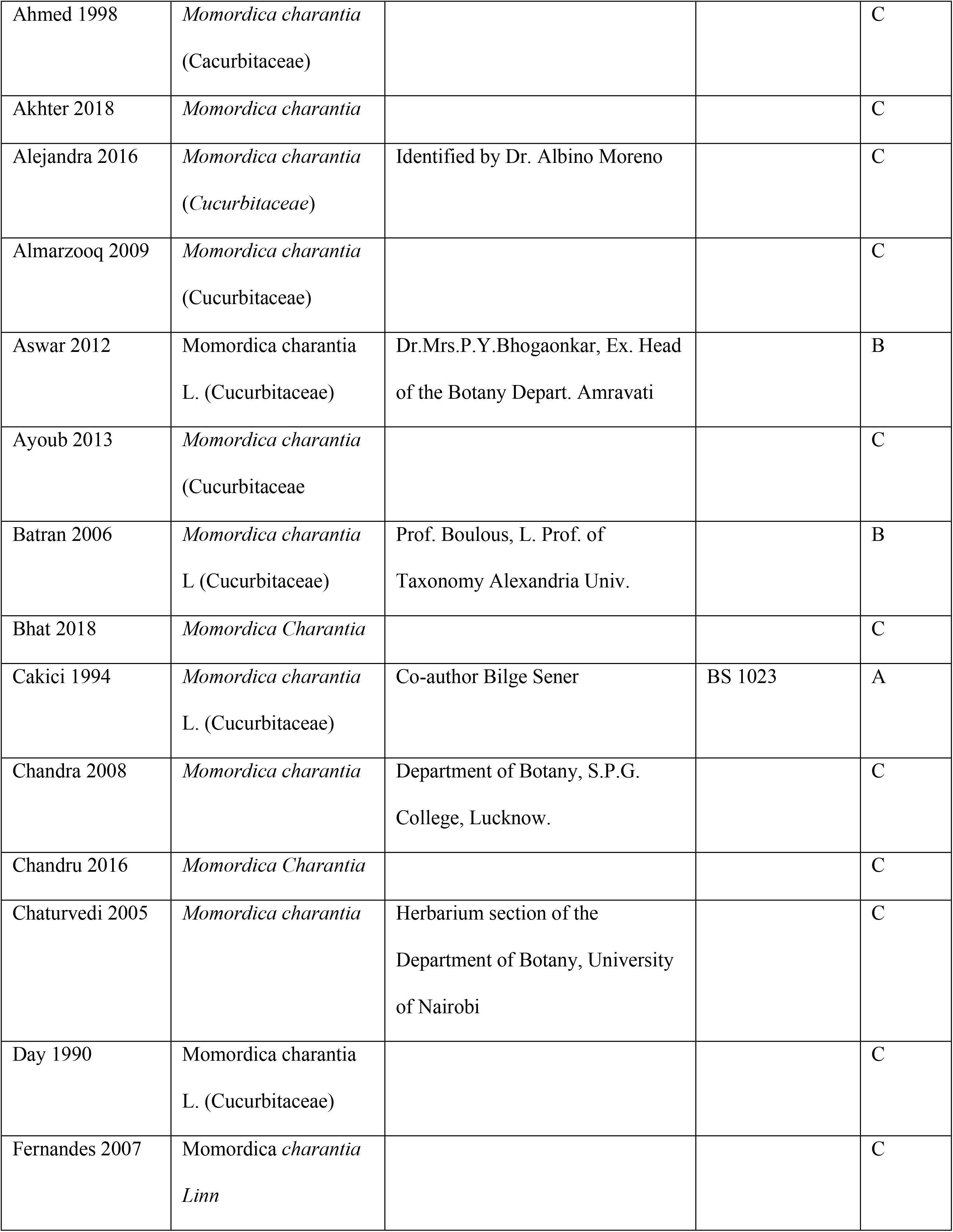
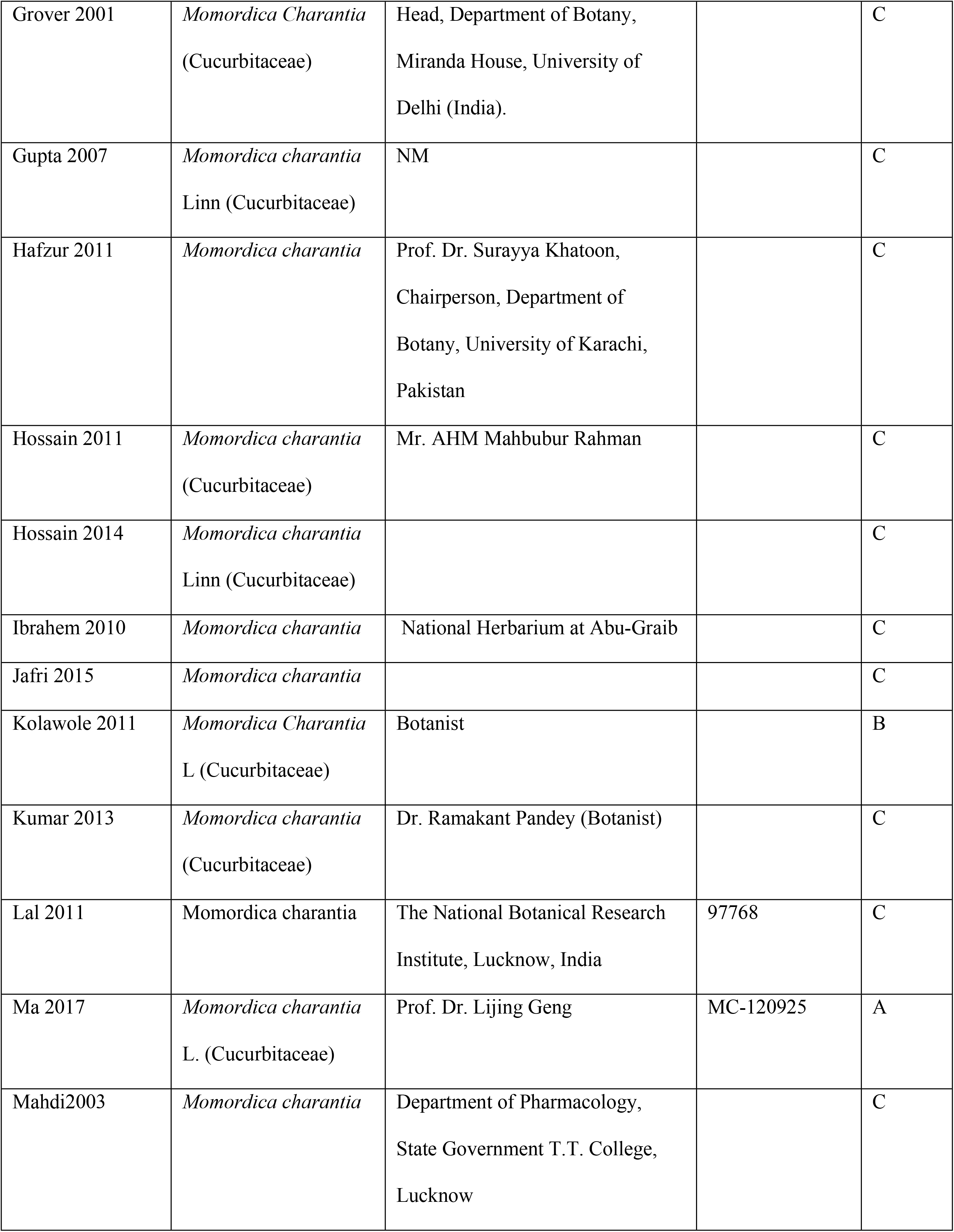
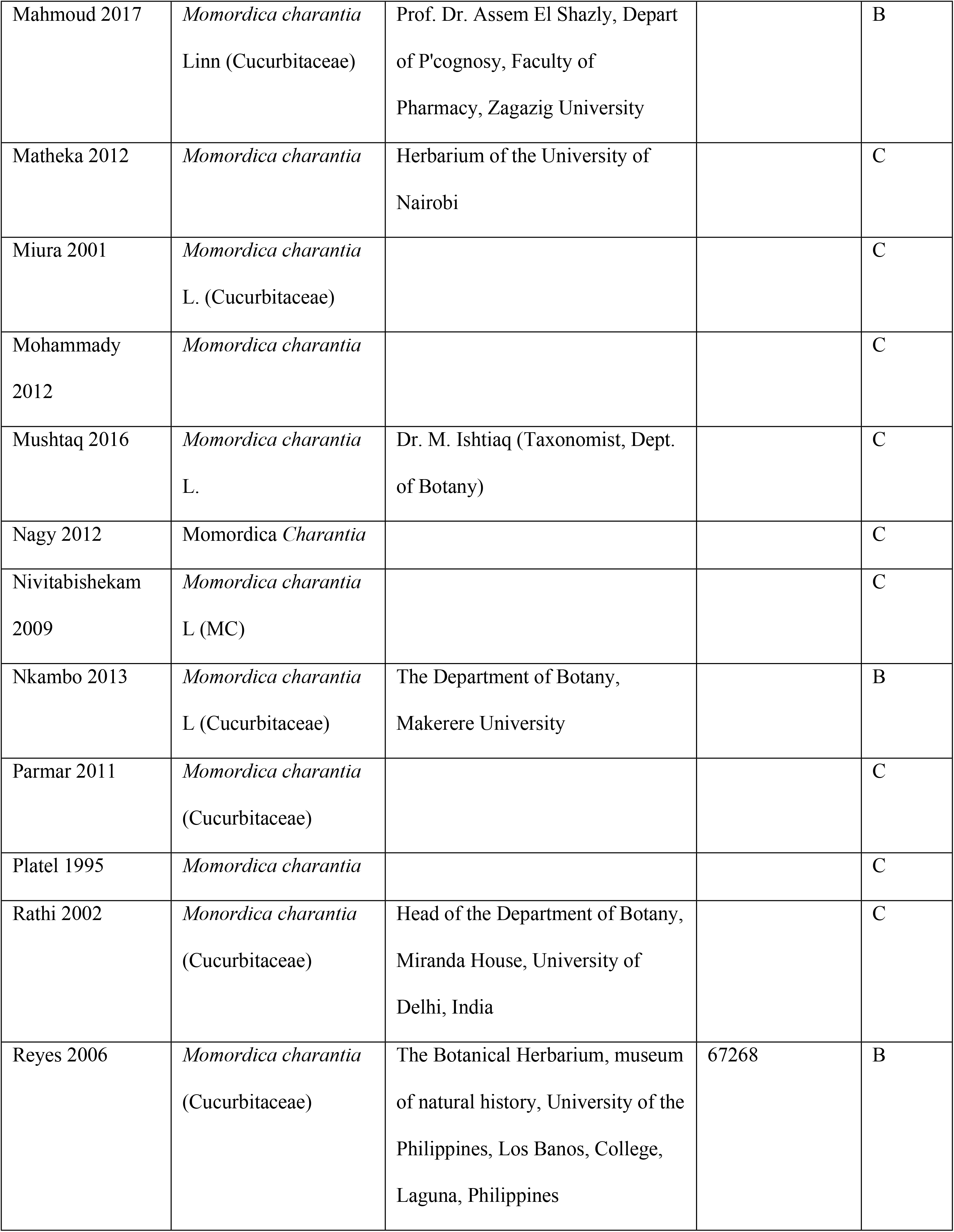
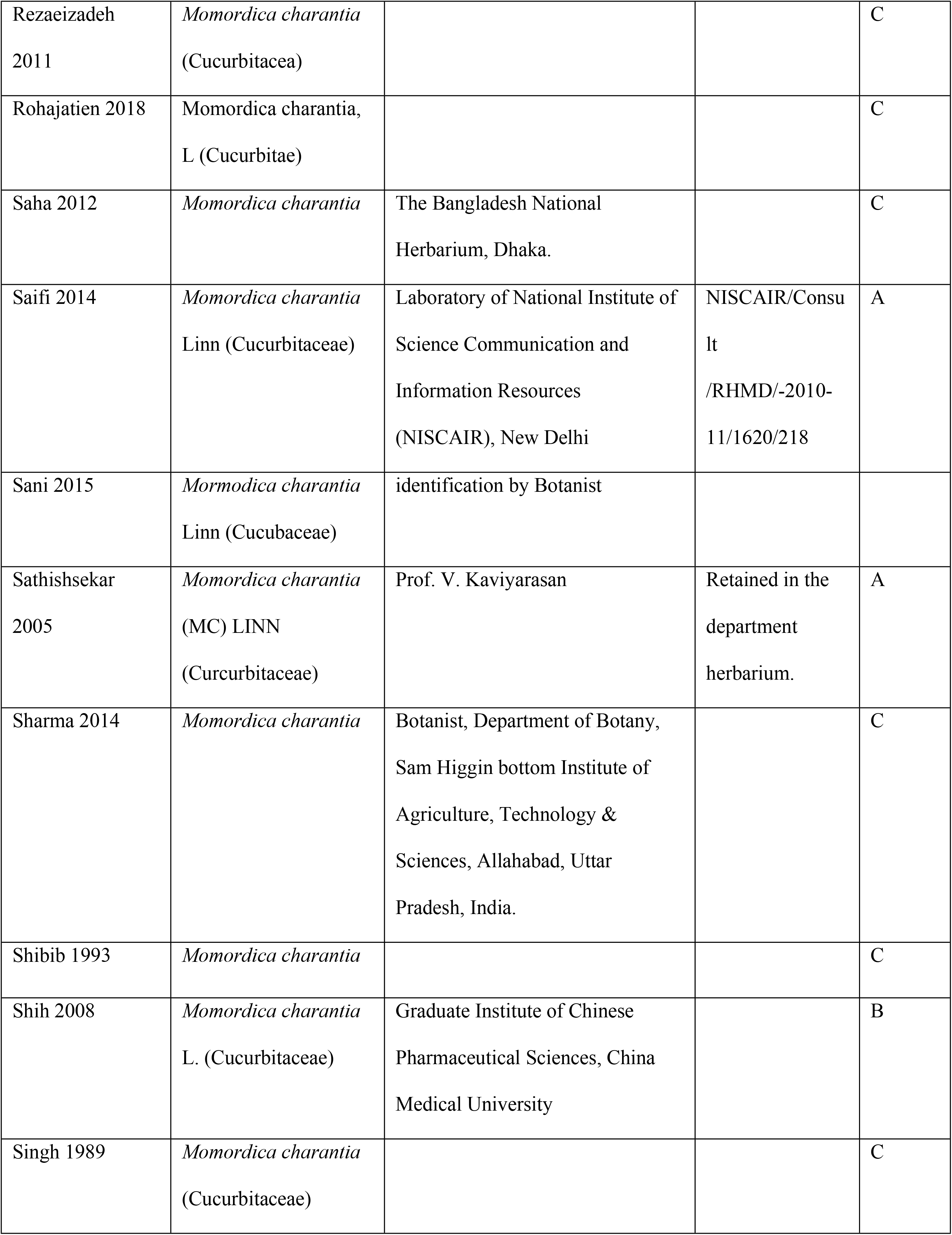
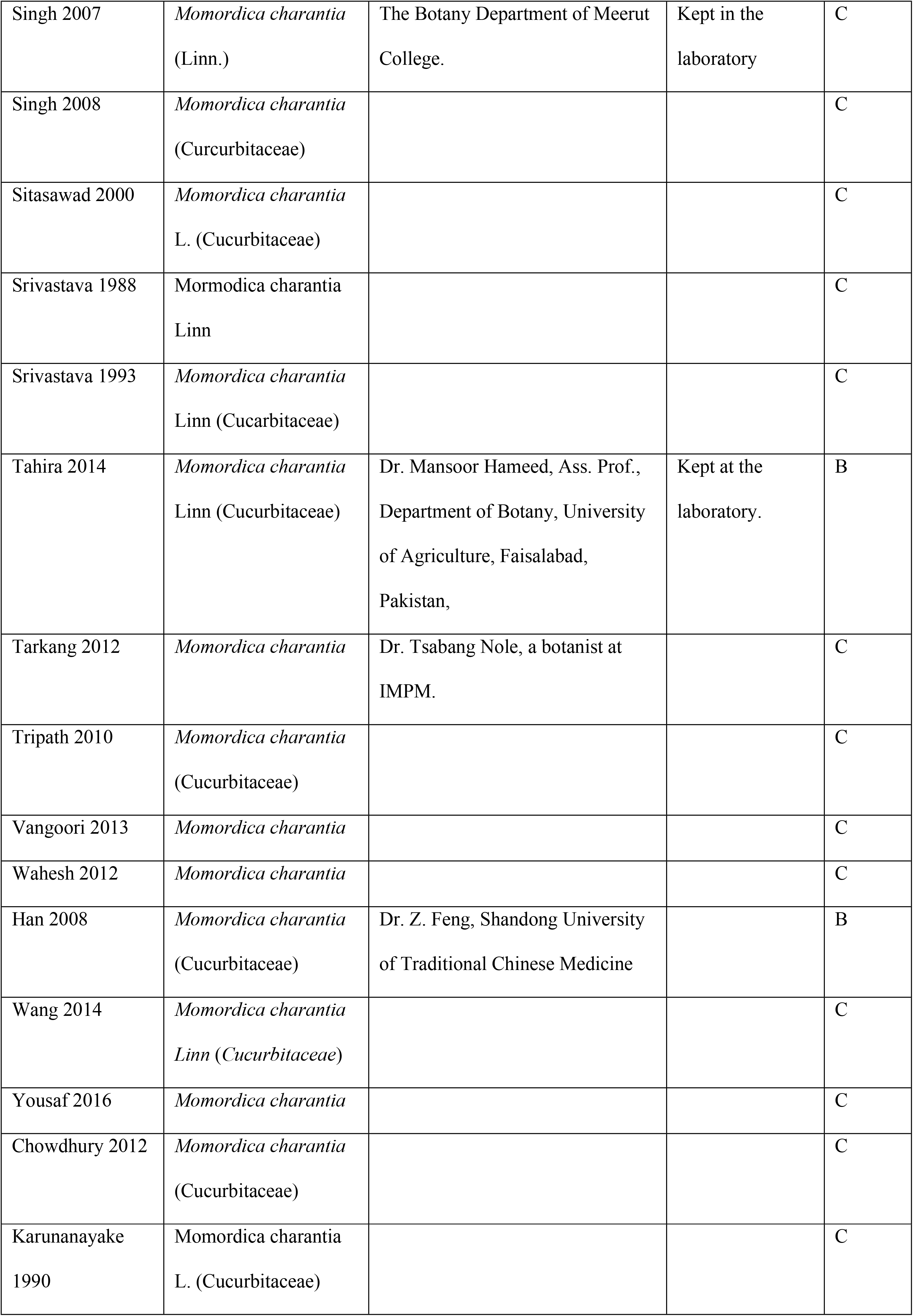
Taxonomical validation score.

### Methodological quality

The quality score of the majority of studies included in the analysis 51 (77.3%) was between 2 and The median score was 3 (interquartile range 1), which means that these studies had poor methodological quality. Interestingly, all 66 studies reported publication in a peer-reviewed journal and claimed to have used appropriate animal models of T2DM. However, none of these studies described the method of random allocation of animals to the treatment or control group, blinded caregiver/investigator, blinded assessment of outcome, and sample size calculation. Surprisingly, only one study described co-interventions of animal models used at baseline [66]. About 25 studies reported compliance with animal welfare regulations while 21 studies reported a statement of maintaining a constant temperature, and only 14 studies provided a statement of potential conflict of interest. Table 5 Summarizes the methodological quality assessment of studies included in the analysis.

**Table 5.** Methodological quality assessment of the included studies.

| Study ID | 1 | 2 | 3 | 4 | 5 | 6 | 7 | 8 | 9 | 10 | Score | Quality Grade |
| --- | --- | --- | --- | --- | --- | --- | --- | --- | --- | --- | --- | --- |
| Abas 2014 | X |  |  |  |  |  | X |  | X | X | 4 | Low |
| Abdollahi 2011 | X | X |  |  |  |  | X |  | X |  | 4 | Low |
| Ahmed 2004 | X |  |  |  |  |  | X |  |  |  | 2 | Low |
| Ahmed 1998 | X |  |  |  |  |  | X |  |  |  | 2 | Low |
| Akhter 2018 | X |  |  |  |  |  | X |  |  | X | 3 | Low |
| Alejandra 2016 | X |  |  |  |  |  | X |  | X |  | 3 | Low |
| Almarzooq 2009 | X | X |  |  |  |  | X |  |  |  | 3 | Low |
| Aswar 2011 | X |  |  |  |  |  | X |  | X |  | 3 | Low |
| Ayoub 2013 | X |  |  |  |  |  | X |  | X |  | 3 | Low |
| Batran 2006 | X |  |  |  |  |  | X |  | X |  | 3 | Low |
| Bhat 2018 | X | X |  |  |  |  | X |  | X | X | 5 | Low |
| Cakici 1994 | X |  |  |  |  |  | X |  |  |  | 2 | Low |
| Chandra 2008 | X |  |  |  |  |  | X |  |  | X | 3 | Low |
| Chandru 2016 | X | X |  |  |  |  | X |  | X | X | 5 | Low |
| Chaturvedi 2005 | X |  |  |  |  |  | X |  |  |  | 2 | Low |
| Day 1990 | X | X |  |  |  |  | X |  |  |  | 3 | Low |
| Fernandes 2007 | X | X |  |  |  |  | X |  | X | X | 5 | Low |
| Grover 2001 | X |  |  |  |  |  | X |  |  |  | 2 | Low |
| Gupta 2007 | X | X |  |  |  |  | X |  |  |  | 3 | Low |
| Hafzur 2011 | X |  |  |  |  |  | X |  |  |  | 2 | Low |
| Hossain 2011 | X | X |  |  |  |  | X |  |  |  | 3 | Low |
| Hossain 2014 | X | X |  |  |  | X | X |  | X | X | 6 | High |
| Ibrahem 2010 | X |  |  |  |  |  | X |  |  |  | 2 | Low |
| Jafri 2015 | X |  |  |  |  |  | X |  |  |  | 2 | Low |
| Kolawole 2011 | X |  |  |  |  |  | X |  | X |  | 3 | Low |
| Kumar 2013 | X | X |  |  |  |  | X |  |  |  | 3 | Low |
| Lal 2011 | X |  |  |  |  |  | X |  |  |  | 2 | Low |
| Ma 2017 | X |  |  |  |  |  | X |  |  | X | 3 | Low |
| Mahdi 2003 | X | X |  |  |  |  | X |  |  |  | 3 | Low |
| Mahmoud 2017 | X |  |  |  |  |  | X |  |  | X | 3 | Low |
| Matheka 2012 | X |  |  |  |  |  | X |  |  | X | 3 | Low |
| Miura 2001 | X | X |  |  |  |  | X |  |  |  | 3 | Low |
| Mohammady 2012 | X | X |  |  |  |  | X |  |  |  | 3 | Low |
| Mushtaq 2016 | X |  |  |  |  |  | X |  | X |  | 3 | Low |
| Nagy 2012 | X | X |  |  |  |  | X |  | X | X | 5 | Low |
| Nivitabishekam<br>2009 | X |  |  |  |  |  | X |  | X | X | 4 | Low |
| Nkambo 2013 | X |  |  |  |  |  | X |  |  |  | 2 | Low |
| Parmar 2011 | X |  |  |  |  |  | X |  | X |  | 3 | Low |
| Platel 1995 | X |  |  |  |  |  | X |  |  |  | 2 | Low |
| Rathi 2002 | X |  |  |  |  |  | X |  |  |  | 2 | Low |
| Reyes 2006 | X |  |  |  |  |  | X |  |  |  | 2 | Low |
| Rezaeizadeh 2011 | X | X |  |  |  |  | X |  | X |  | 4 | Low |
| Rohajation 2018 | X |  |  |  |  |  | X |  |  |  | 2 | Low |
| Saha 2012 | X | X |  |  |  |  | X |  | X |  | 4 | Low |
| Saifi 2014 | X | X |  |  |  |  | X |  | X |  | 4 | Low |
| Sani 2015 | X |  |  |  |  |  | X |  |  |  | 2 | Low |
| Sathishsekar 2005 | X |  |  |  |  |  | X |  | X |  | 3 | Low |
| Sharma 2014 | X |  |  |  |  |  | X |  | X | X | 4 | Low |
| Shibib 1993 | X |  |  |  |  |  | X |  |  |  | 2 | Low |
| Shih 2008 | X | X |  |  |  |  | X |  | X |  | 4 | Low |
| Singh 1989 | X |  |  |  |  |  | X |  |  |  | 2 | Low |
| Singh 2007 | X |  |  |  |  |  | X |  |  |  | 2 | Low |
| Singh 2008 | X |  |  |  |  |  | X |  | X |  | 3 | Low |
| Sitasawad 2000 | X |  |  |  |  |  | X |  |  |  | 2 | Low |
| Srivastava 1988 | X | X |  |  |  |  | X |  |  |  | 3 | Low |
| Srivastava 1993 | X |  |  |  |  |  | X |  |  |  | 2 | Low |
| Tahira 2014 | X |  |  |  |  |  | X |  | X |  | 3 | Low |
| Tarkang 2012 | X |  |  |  |  |  | X |  |  |  | 2 | Low |
| Tripath 2010 | X | X |  |  |  |  | X |  | X |  | 4 | Low |
| Vangoori 2013 | X |  |  |  |  |  | X |  |  |  | 2 | Low |
| Wahesh 2012 | X |  |  |  |  |  | X |  |  |  | 2 | Low |
| Han 2008 | X |  |  |  |  |  | X |  |  |  | 2 | Low |
| Wang 2014 | X | X |  |  |  |  | X |  | X | X | 5 | Low |
| Yousaf 2016 | X |  |  |  |  |  | X |  |  |  | 2 | Low |
| Chowdhury 2012 | X |  |  |  |  |  | X |  |  |  | 2 | Low |
| Karunanayake<br>1990 | X |  |  |  |  |  | X |  |  |  | 2 | Low |

### Risk of bias assessment

The risk of bias of the preclinical studies included in the analysis was assessed using SYRCLE’s risk of bias tool. The results indicated that all studies included did not perform allocation concealment, random animal housing, blinding of animal caregivers and investigators, random outcome assessment and blinding of outcome assessment. This could mean that these studies were prone to systematic errors due to the design flaw that could overestimate the effect of the *M. charantia* L. Four studies were given unclear risk of bias with regard to random sequence generation because review authors found inadequate description of the method used for random sequence generation [15,56,85,92]. Summary of the risk of bias across all studies and risk of bias of each included study is provided in Fig 2 and S3 appendix respectively.

**Fig 2.**
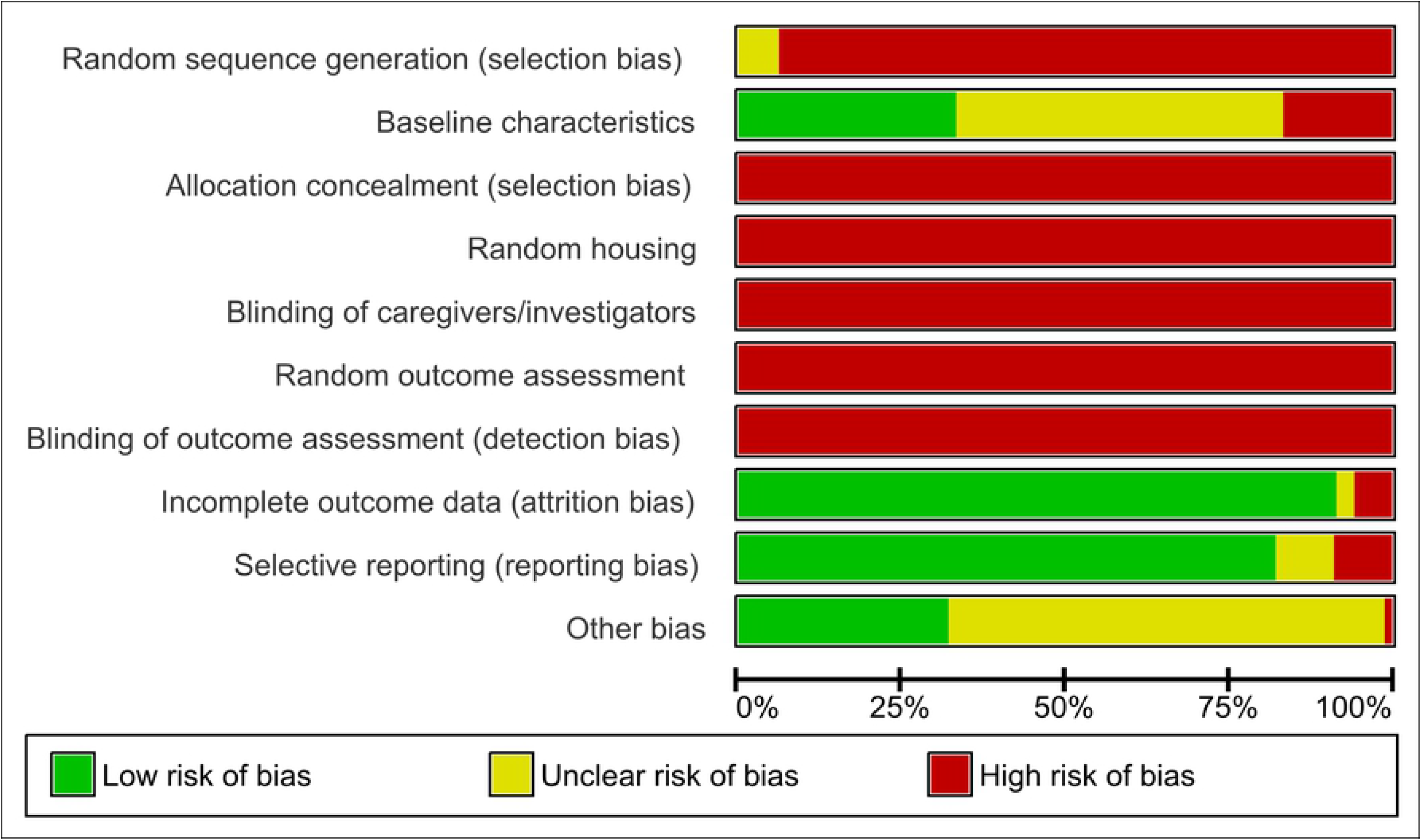
Risk of bias graph: review authors’ judgements about each risk of bias item presented as percentages across all included studies.

### Effect of intervention

#### Primary outcome; Fasting Plasma Glucose (FPG)

About 42 preclinical studies (n = 815) had data on FPG. The pooled estimate indicated moderate quality evidence that *M. charantia* L. significantly reduced FPG compared with a vehicle control group, representing −6.86 of SMD (95% CI; −7.95, −5.77), I^2^ = 90. Interestingly, all studies consistently favored *M. charantia* L. (Fig 3).

**Fig 3.**
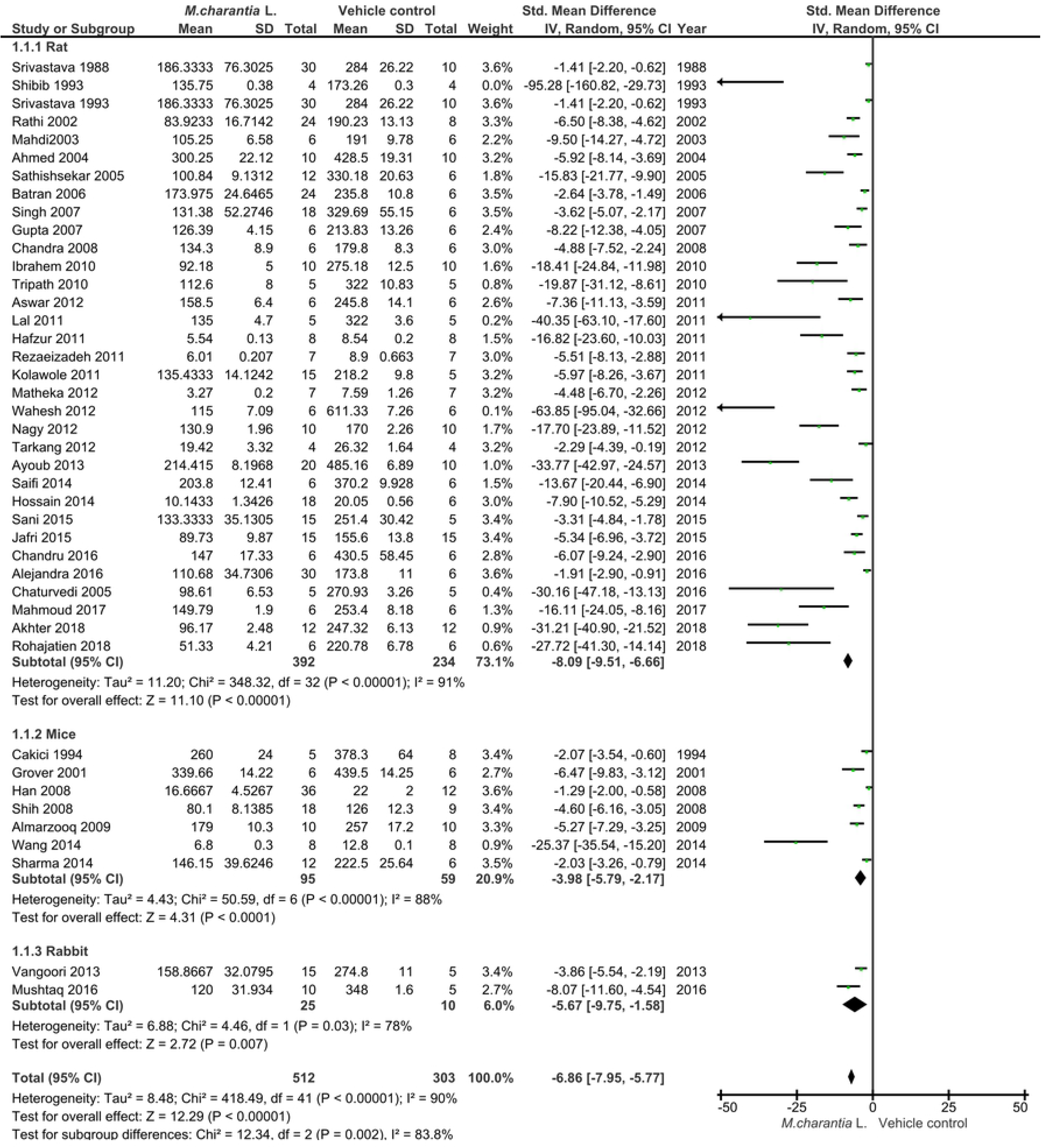
Forest plot of preclinical studies comparing *M. charantia* L. and vehicle control; measuring fasting plasma glucose

#### Secondary outcomes

##### a. Glycosylated hemoglobin A1c (HbA1c)

The data from three preclinical studies were pooled for assessment of HbA1c (Fig 4). There was moderate quality evidence that *M. charantia* L. significantly lowered HbA1c level in a treated group (n = 34) compared to the vehicle control group (n = 25); −7.76 of SMD (95% CI; −12.5, - 3.01). The I^2^ = 82% was indicating the presence of heterogeneity in individual studies.

**Fig 4.**
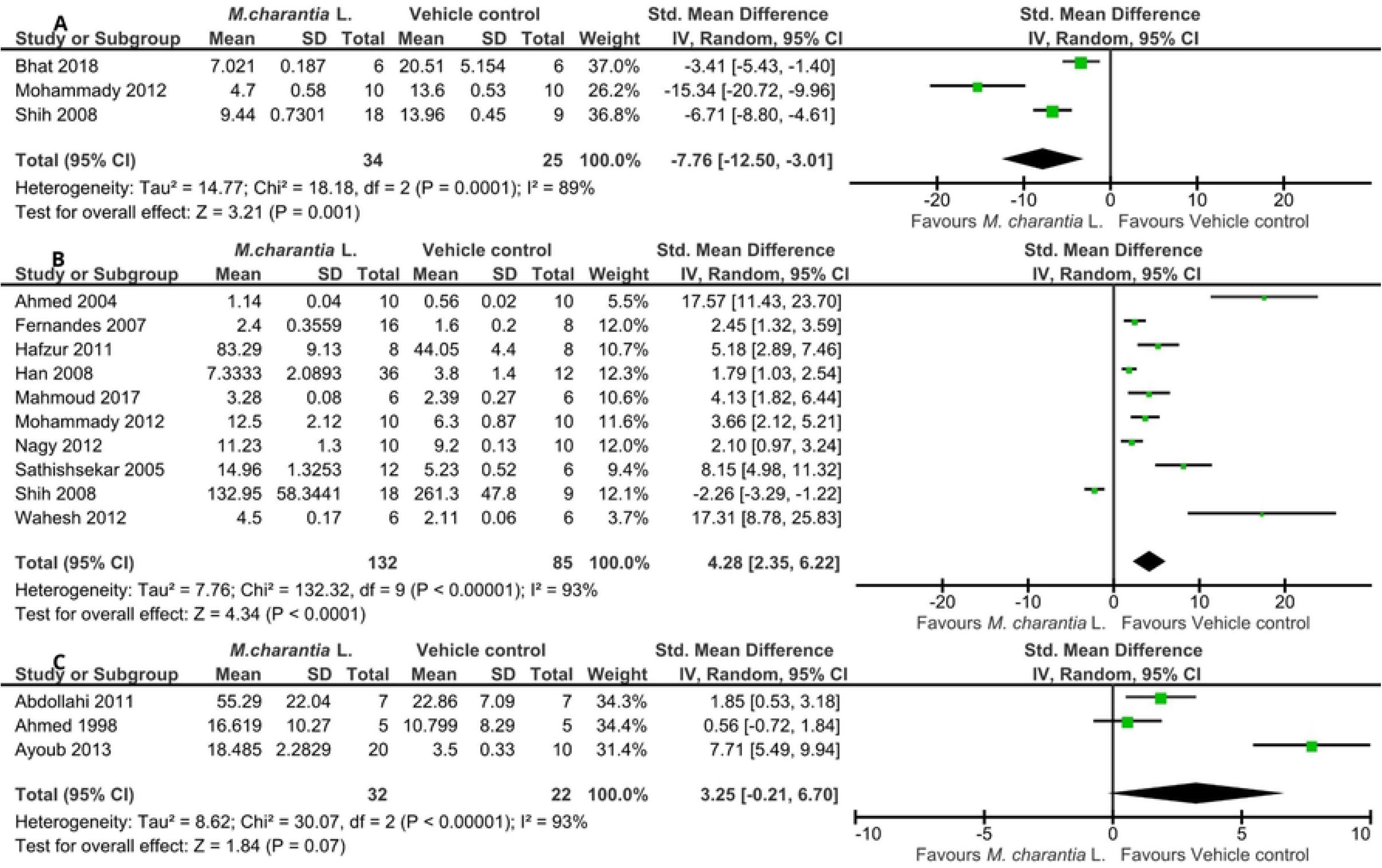
Forest plot of preclinical studies comparing *M. charantia* L. and vehicle control; measuring (A) HBA1c, (B) Serum insulin level, (C) Insulin-positive cells in pancreases

##### b. Serum insulin

Results indicated very low-quality evidence that serum insulin level observed in *M. charantia* L. treated group (n = 132) was significantly increased compared with the vehicle control group (n = 85); 4.28 of SMD (95% CI; 2.35, 6.22). The I^2^ of 93% was indicating the presence of heterogeneity. Only one study (Shih et al., 2008) had a significant effect size in the opposite direction (Fig. 4). The review authors downgraded the evidence to very low-quality due to a severe risk of bias and imprecision, serious inconsistency, and strongly suspected publication bias (S4 Appendix).

##### c. Insulin-positive cells

There was an increase in the number of insulin-positive cells in the *M. charantia* L. treated group (n = 32) compared to the vehicle control group (n = 22); 3.25 of SDM (95% CI; −0.21, 6.70). Although such an increase was not statistically significant, all the three studies favored the intervention (Fig. 4). The I^2^ was 93% indicated the presence of heterogeneity in the individual study.

##### d. Liver glycogen

The pooled data from four studies indicated that there was no statistically significant increase in liver glycogen in the *M. charantia* L. treated group (n = 56) compared to the vehicle control group (n = 36); 1.11 of SDM (95% CI; −3.20, 5.43). The I^2^ of 97% indicated the presence of heterogeneity. Two studies favored *M. charantia* L., and another two studies favored the vehicle control group (Fig 5).

**Fig 5.**
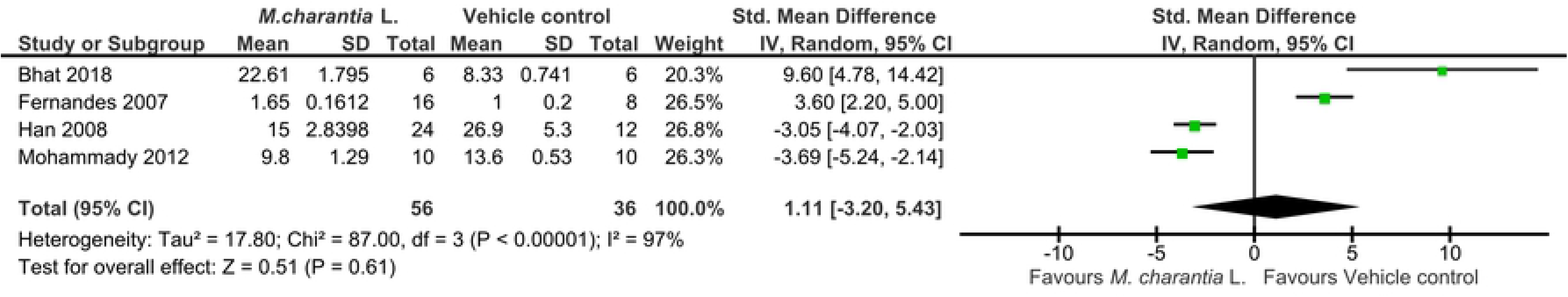
Forest plot of pre-clinical studies comparing *M. charantia* L. and vehicle control; measuring liver glycogen

##### e. Triglycerides (TGs)

The data from 13 preclinical studies were pooled for assessment of triglycerides (Fig 6). Results showed a very low-quality evidence that *M. charantia* L. significantly lowered TGs level in treated group (n = 142) compared to vehicle control group (n = 87); −9.12 of SMD (95% CI; −11.76, - 6.49). The I^2^ was 92% indicated the presence of substantial heterogeneity in individual studies.

**Fig 6.**
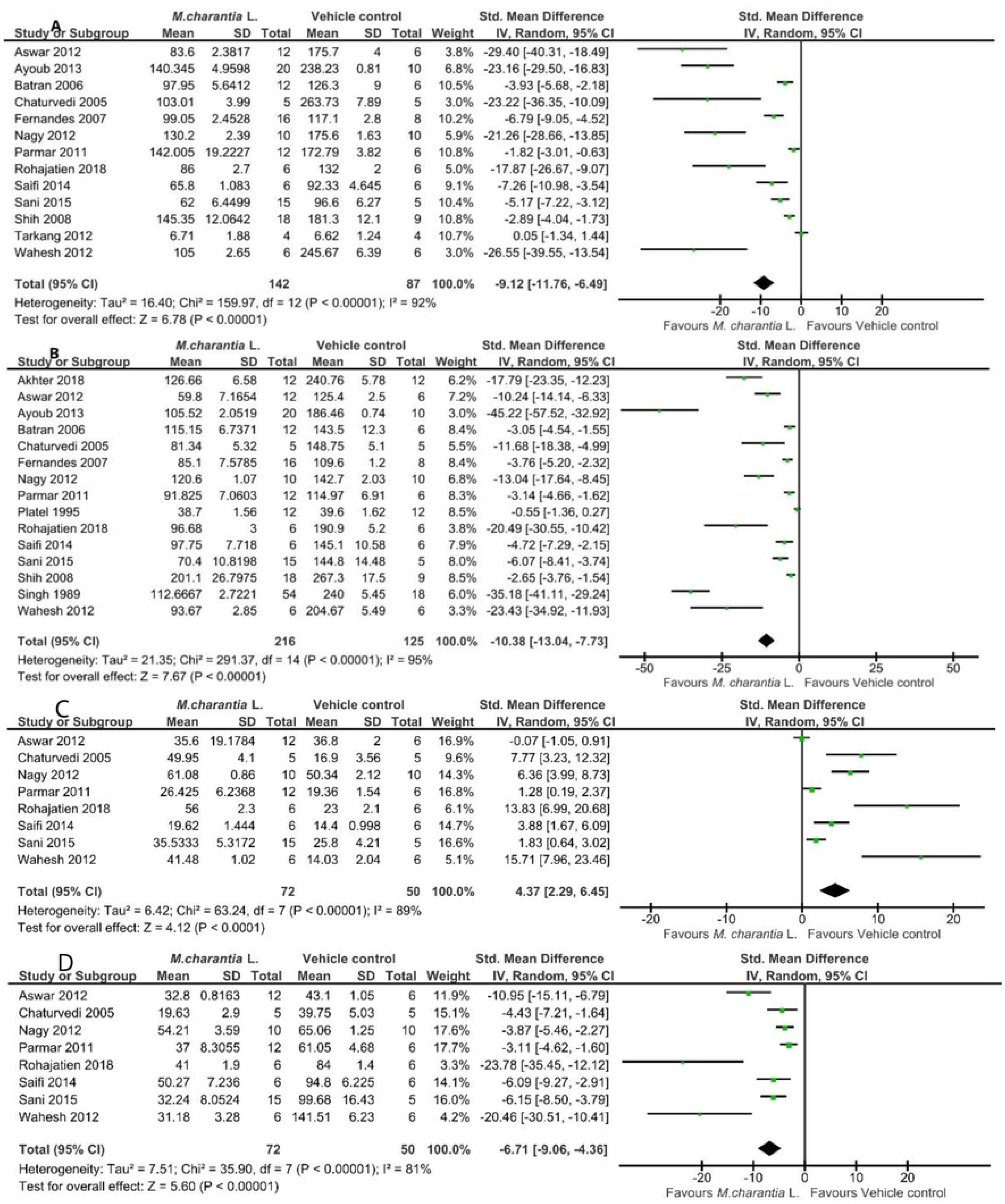
Forest plot of preclinical studies comparing *M. charantia* L. and vehicle control; measuring (A) TGs, (B) TC, (C) HDL-c, (D) LDL-c

##### f. Total cholesterol (TC)

Figure 6 showed that *M. charantia* L. treated group (n = 216) had a significantly reduced level of total cholesterol compared with the vehicle control group (n = 125) with SMD of −10.38 (95% CI; −13.04, −7.73). The I^2^ was 95% indicated the heterogeneity. The certainty of this evidence was assessed as low (S4 Appendix).

##### g. High-density lipoprotein cholesterol (HDL-c)

The HDL-c was assessed by integrating data from eight studies (Fig 6). There was low quality-evidence that the HDL-c level in *M. charantia* L. treated group (n = 72) increased compared to the vehicle control group (n = 50), 4.37 SDM (95% CI; 2.29, 6.45). The I^2^ was 89% indicated the presence of heterogeneity.

##### h. Low-density lipoprotein cholesterol (LDL-c)

The LDL-c level in the *M. charantia* L. treated group (n = 72) was significantly decreased compared to that observed in the vehicle control group (n = 50). The SMD of −6.71 (95% CI; - 9.06, −4.36). The I^2^ was 89% indicated the presence of heterogeneity (Fig 6).

##### i. Alanine aminotransferase (ALT), Aspartate aminotransferase (AST), and Alkaline phosphate (ALP)

There was a significant reduction of ALT (SMD; −5.14; [95% CI; −7.33, −2.95]), AST (SMD; - 3.60; [95% CI; −4.95, −2.25]), and ALP (SMD; −3.58; [95% CI; −4.94, −2.22]) in the *M. charantia* L. treated groups compared with the vehicle control groups. The I^2^ were 77%, 64% and 82% respectively; indicated the presence of significant heterogeneity (Fig 7).

**Fig 7.**
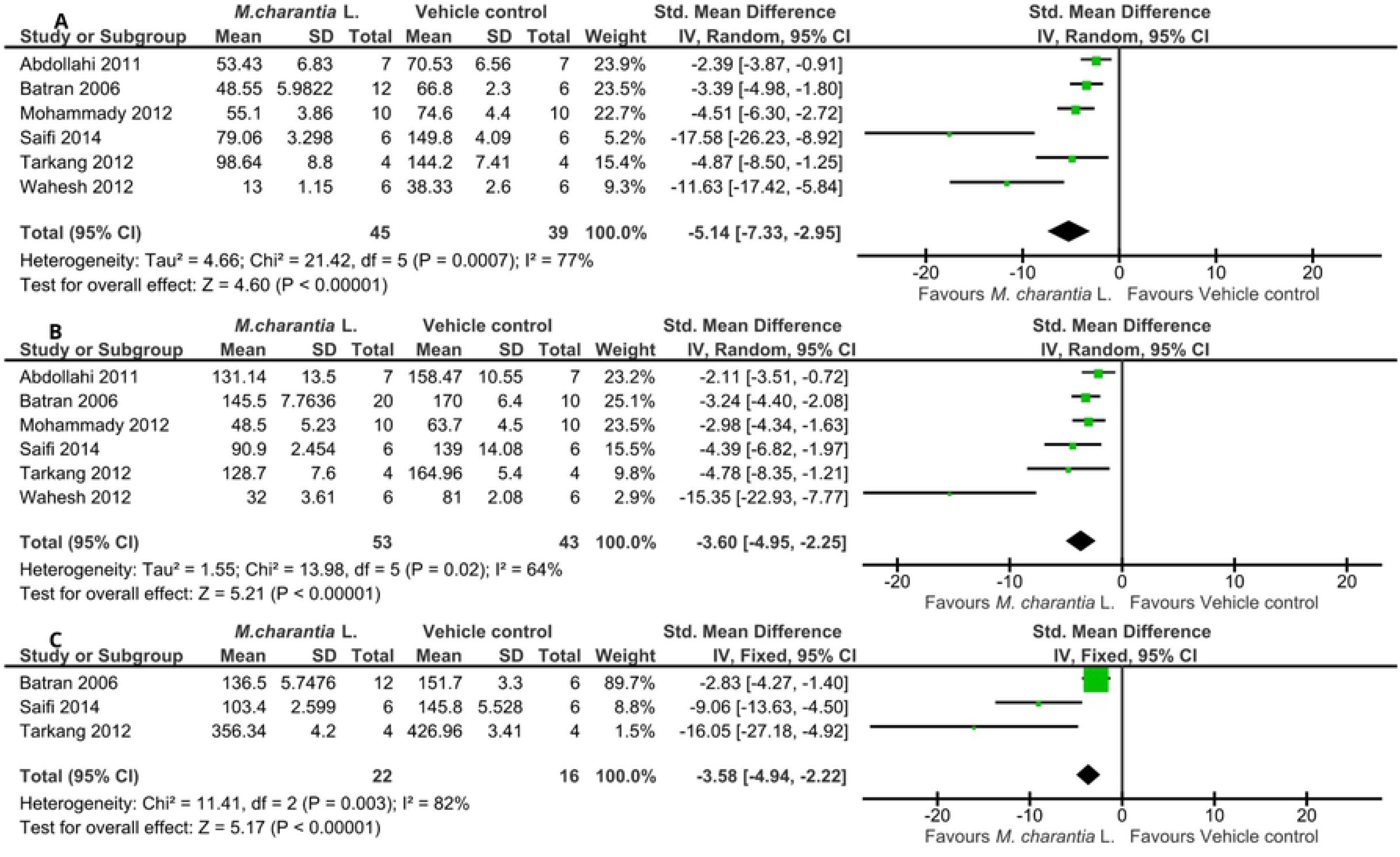
Forest plot of preclinical studies comparing *M. charantia* L. and vehicle control; measuring (A) ALT, (B) AST, (C) ALP

##### j. Serum creatinine and plasma urea

There was a significant reduction of serum creatinine (SMD; −4.52; [95% CI; −6.42, −2.61]) and plasma urea (SMD; −2.68; [95% CI; −4.13, −1.22]) in the *M. charantia* L. treated groups compared with the vehicle control groups. The I^2^ were 84% and 89% respectively; indicated the presence of significant heterogeneity (Fig 8).

**Fig 8.**
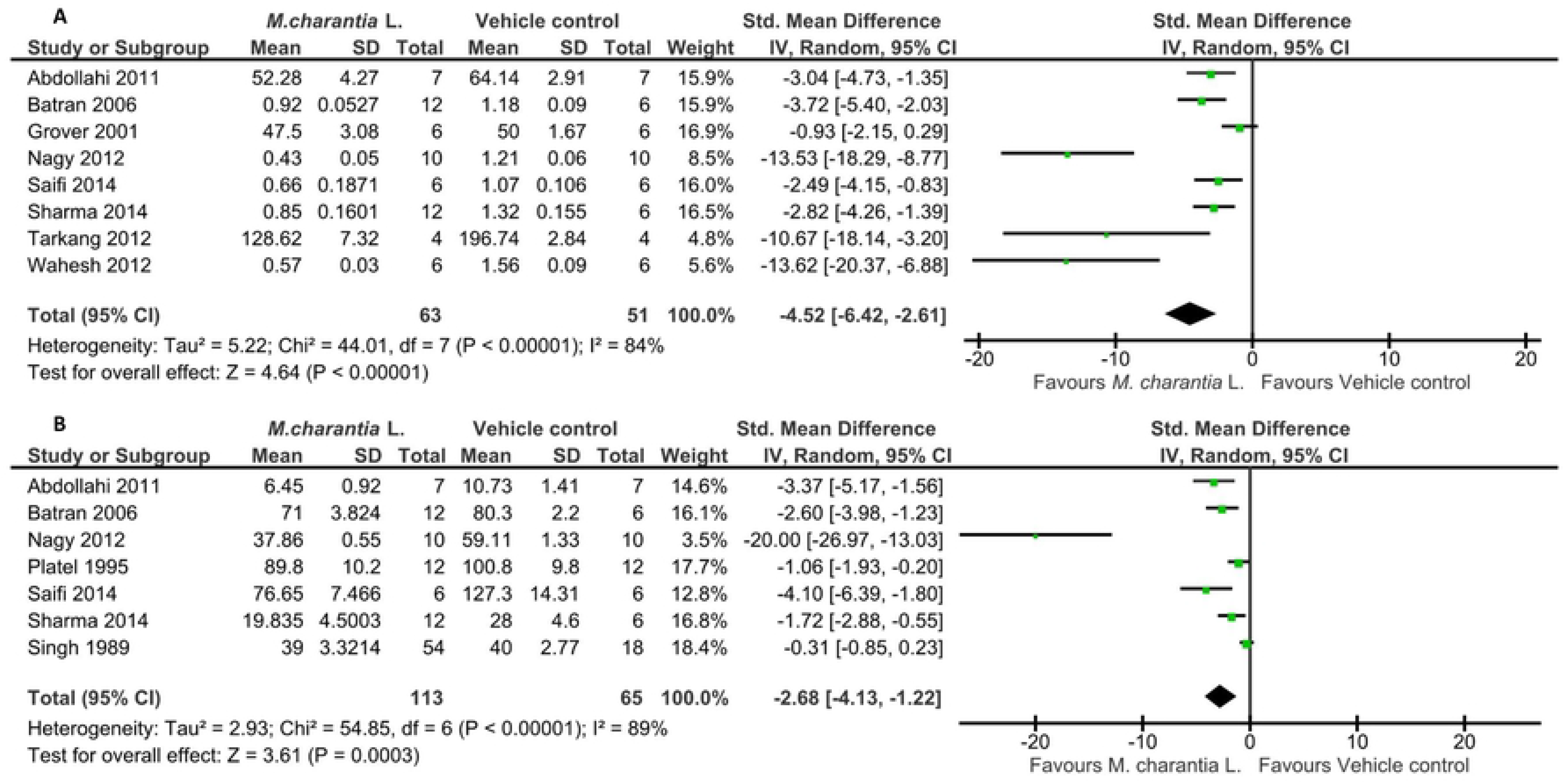
Forest plot of preclinical studies comparing *M. charantia* L. and vehicle control; measuring (A) Serum creatinine, (B) Plasma urea

##### k. Effect of M. charantia L. on body weight

About 14 preclinical studies provided quantitative data on weight. We reported a significant increase in body weight in the *M. charantia* L. treated groups (n = 148) compared with the vehicle control groups (n = 114). The SMD was 2.96 (95% CI; 1.63, 4.29), and I^2^ was 56% which indicates the presence of moderate heterogeneity in individual studies (Fig 9).

**Fig 9.**
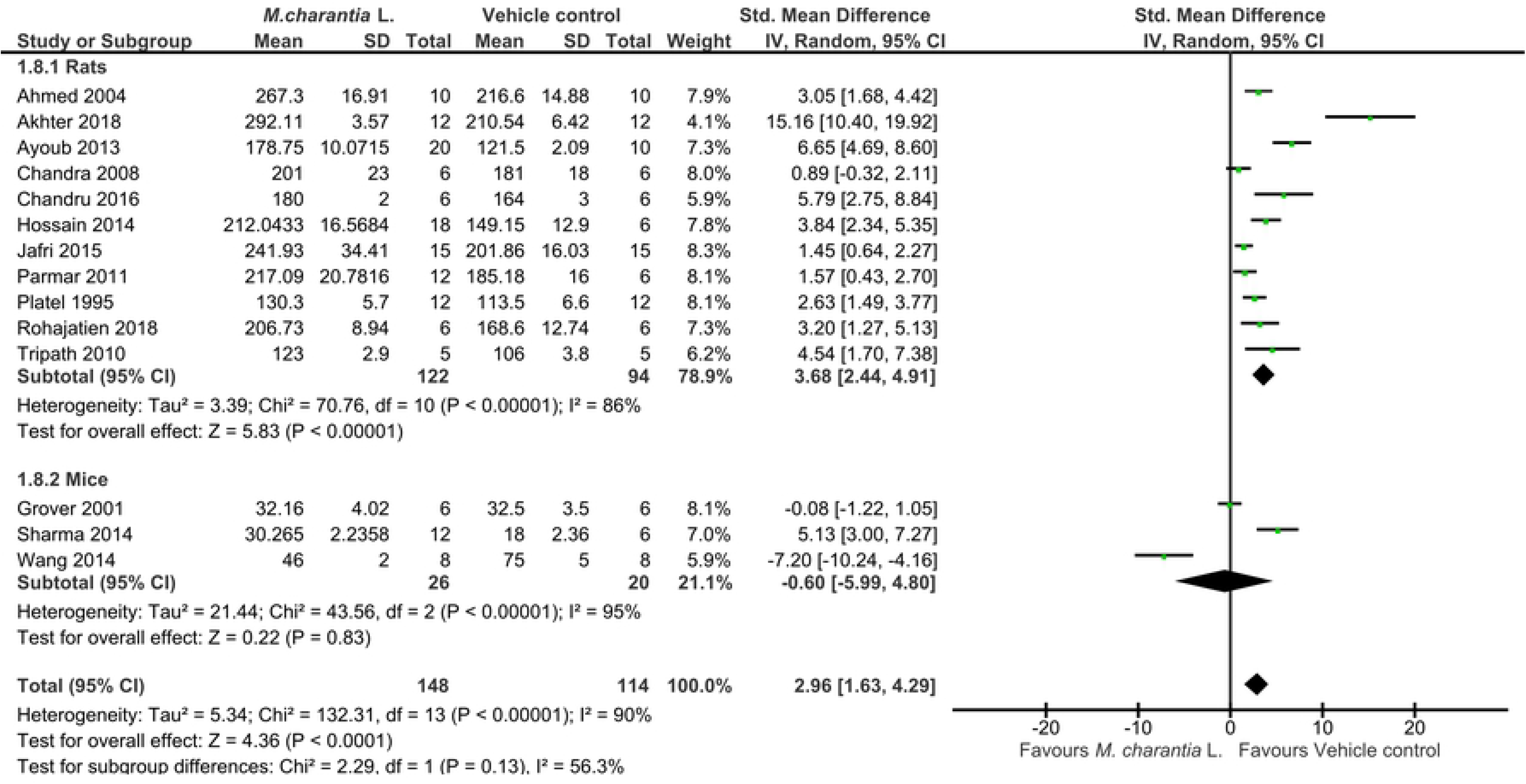
Forest plot of preclinical studies comparing *M. charantia* L. and vehicle control; measuring body weight

### Subgroup analysis

Review authors considered subgroup analysis for *M. charantia* L. versus vehicle control on FPG level for rats, mice and rabbits’ species (Fig. 4). The test for subgroup analysis suggested that there was a statistically significant subgroup effect (P = 0.002), I^2^ = 83.8%, meaning that animal species significantly modified the effect of *M. charantia* L. in comparison to vehicle control. The treatment effect favored *M. charantia* L. across all the three species. However, the treatment effect was higher for rats than mice and rabbits. Hence the subgroup effect is quantitative. However, there was still heterogeneity between results in studies within subgroup which requires further exploration.

Another subgroup analysis considered for *M. charantia* L. versus vehicle control on FPG level was animal strain; Wistar albino rats, Sprague-Dawley rats, and Charles Foster rats. The test for subgroup differences indicated that there was a statistically significant subgroup effect (P = <0.00001), I^2^ = 85.4%, meaning that animal strains significantly modified the effect of *M. charantia* L. in comparison to vehicle control. The treatment effect favored *M. charantia* L. over vehicle control for all animal strains (Wistar albino rats, Sprague-Dawley rats, and Charles Foster rats); therefore, the subgroup effect is quantitative. It is interesting to note that effect size was greater for Wistar albino rats (SMD; −10.29, 95%CI; −12.55, −8.03), I^2^ = 90% than for Sprague-Dawley rats (SMD; −6.71, 95%CI; −10.02, −3.40), I^2^ = 88%, and Charles Foster rat (SMD; −2.15, 95%CI; −3.69, −0.60), I^2^ = 80%. This analysis indicated a substantial unexplained heterogeneity between the studies with each subgroup as indicated by their I^2^. It is also important to note that the subgroup analysis could not detect subgroup effects of other animal stains (Horts men rats, Long-Evans rats, Swiss albino mice, Kunming mice, C57BL/6J and KK/HIS mice) because a small number of studies contributed the data.

### Evaluation of publication bias

Review authors assessed publication bias for *M. charantia* L. versus vehicle control on FPG, TC, TGs, serum insulin and body weight by visually assessing funnel plots. Egger’s tests for funnel plot asymmetry suggested that there was a statistically significant publication bias for FPG, TC, TGs, and serum insulin (P <0.0001). However, there was no evidence of publication bias for the weight (P = 0.062).

The trim and fill analysis imputed 19 potentially missed experimental studies for FPG, 9 for TC, 5 for TGs and one missed study for serum insulin. The imputed missed experimental studies changed the significance or magnitude of the overall pooled effect size for these outcomes. Use of random effect models indicated that FPG changed to non-significant reduction −2.46 SMD, (95%CI; −5.10, 0.17), similarly, the TGs changed to −1.95 SMD, (95%CI; −8.67, 4.76), while the magnitude of effect size of TC reduced to −2.22 SMD (95%CI; −2.68, −1.77) from the original analysis, and that of serum insulin increased to 4.58 SMD (95%CI; 0.65, 8.50). The fig 10A and 10B illustrate the effect of adjustment by trim and fill analysis.

**Fig10A.**
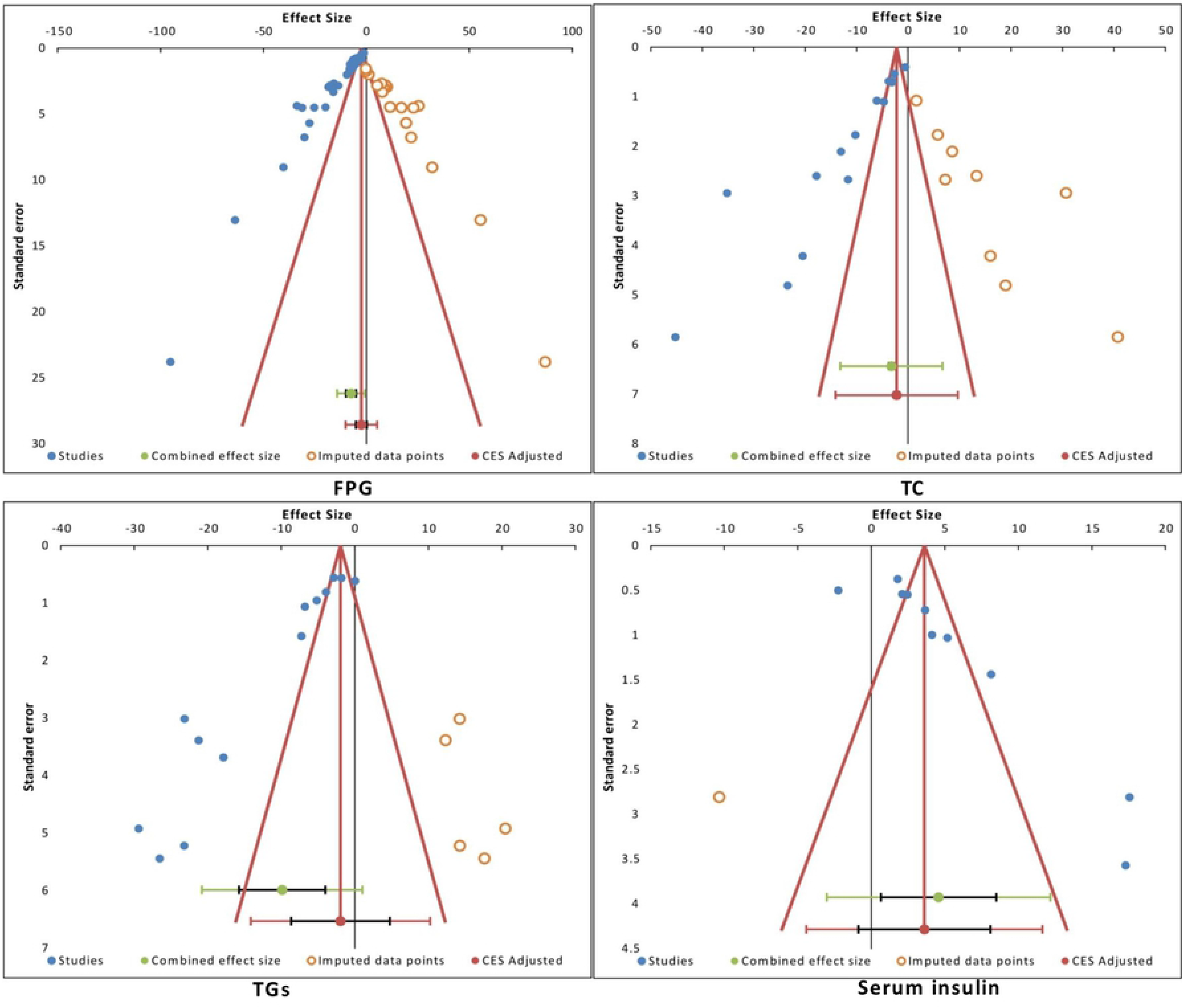
Trim and Fill adjustment of publication bias on the standardized mean difference for fasting plasma glucose (FPG), total cholesterol (TC), triglycerides (TGs), and serum insulin.

**Fig 10B.**
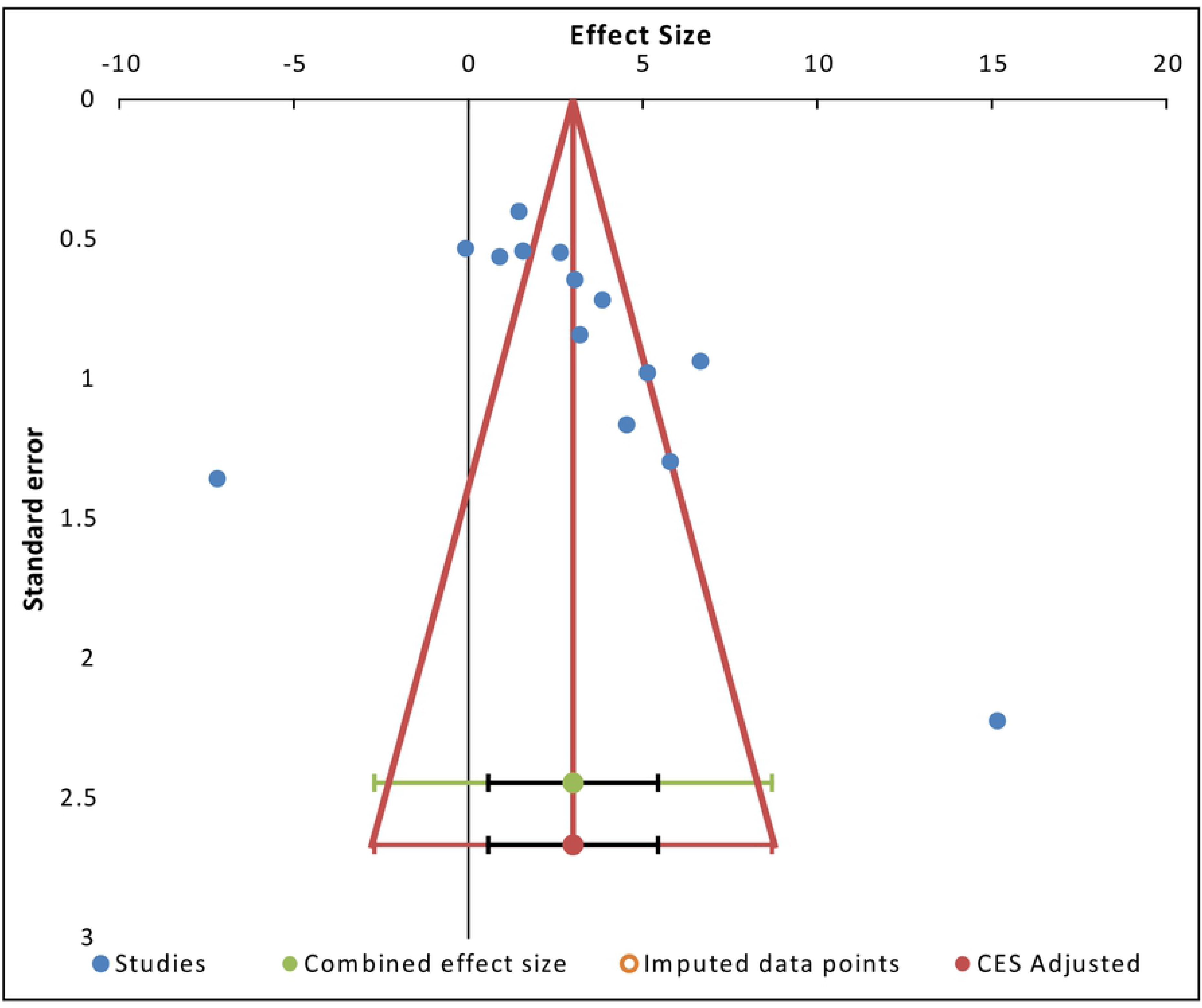
Trim and Fill adjustment of publication bias on the standardized mean difference for body weight

## Discussion

### Summary of the main findings

This systematic review and meta-analysis is to the best of our knowledge the first to provide quantitative estimates of the effect of *M. charantia* L. on essential attributes of type 2 diabetes mellitus in experimental studies. The cumulative evidence concludes that the administration of *M. charantia* L. to animal models of T2DM can reduce fasting plasma glucose level. Review authors grade the quality of evidence as moderate because of the very serious risk of bias of included studies, strong evidence of publication bias and unexplained heterogeneity. The suspected publication bias in our meta-analysis could have impacted on the overall effect of *M. charantia* L. on FPG. About 19 studies deemed missing and adjusted estimate reduced FPG level to non-significant, suggesting that our findings may be inflating *M. charantia* L. efficacy. However, we should consider this interpretation with caution because the authors assessed publication bias only by considering the asymmetric nature of the funnel plot. It should be pointed out that publication bias is not the only explanation for funnel plot asymmetry. Previous studies have established several other factors that could lead to asymmetric funnel plot such as true heterogeneity, poor methodological quality, artefactual, and chance [43]. With these factors become ubiquitous in preclinical studies, as reported in other assessments [106–108], it could be safe to assume their influence on funnel plot asymmetry observed. The finding that publication bias overstated effect size is in line with other systematic reviews studies [109, 110].

The present results of meta-analysis confirmed that administering *M. charantia* L. extracts for at least three months could increase serum insulin level, HDL-c, and body weight while significantly reduced HbA1c, triglycerides, total cholesterol, LDL-c, ALT, AST, ALP, urea, and creatinine. The plausible explanation for the increase in serum insulin level could be that *M. charantia* L. works by enhancing insulin release from the partially destroyed beta-cells in the pancreases or increase beta-cell mass or both. The results of qualitative synthesis supported this argument. Seven studies reported that *M. charantia* L. increases the number of β-cells in the pancreas thereby improving the capability to produce insulin [22,51,56] and partially restore the healthy cellular population and enlarged size of islets with hyperplasia [15,47,55,56,66]. The observed decrease in triglycerides, total cholesterol, LDL-c and increase in HDL-c underscore the potential of *M. charantia* L. in controlling type 2 diabetes mellitus and its associated complications. The review results further confirm the hepato-renal protective effect of *M. charantia* L. that could partly explain the long history of its use as a nutritional food and herbal medicines for various ailments in local communities in Asia, South America, and Africa [111, 112].

### Quality of the evidence

Study methodological quality is a critical factor that threatens the validity of preclinical studies. According to the CAMARADES quality score, all studies included in our meta-analysis are of poor methodological quality with an average score of 3. Besides, the SYRCLEs risk of bias tool assessed all studies as having a high risk of bias in the domains of random sequence generation, allocation concealment, random housing, blinding investigators/caregivers, random outcome assessment, and blinding outcome assessment. High risk of bias in these domains means that the studies have poor internal validity. It is now clear that these aspects of experimental design can have a substantial impact on the reported outcome of experiments [113]. While researchers recognized the importance of these issues for decades, they are rarely reported in publications of animal experiments [108].

Our review found that two studies out of 66 assessed, did not report the number of animals used. While the studies that reported, have no description of the method used to calculate the sample size. Similar findings were reported in another review of animal studies in India [107]. Reporting animal numbers is essential so that the biological and statistical significance of the experimental results can be assessed or the data reanalyzed and is also necessary if the experimental methods are to be repeated [114]. Appropriate sample size calculation ensures a study designed with sufficient power to detect the true effect of the intervention; thus failure to use adequate sample size can potentially have an impact on science, ethics, and economy.

The meta-analysis reported higher heterogeneity because included studies used different methodological design features such as; different induction materials for T2DM, different dose of interventions, duration of administration, different types of extracts, different outcome measurement scales, and the small sample size. Of particular concerns is the high dose of STZ and alloxan used to induce T2DM. Although these induction materials are widely used to induce experimental T2DM, alloxan can cause kidney toxicity due to its very narrow effective dose and a higher dose of STZ could completely knock off beta-cells and potentially induce type 1 diabetes mellitus [115]. The previous study indicated that a single dose of 45mg/kg STZ leads to hyperglycemia and a higher mortality rate than multiple doses of 30mg/kg [116]. Inspired by a growing understanding of disease pathophysiology, researchers have now revealed that a combination of high-fat diet and low dose STZ produce a model of T2DM that closely mimic a natural history of human with T2DM [117, 118]. Our findings suggest that the concern about different model of inducing T2DM varying similarity to human with the condition is warranted. These design features could potentially be sources of heterogeneity, and by extension, influence constructs validity of the study.

The meta-analysis included studies from at least 20 different countries, meaning that *M. charantia* L. can have varying constituents according to the region of origin, harvesting season, mode of cultivation, or different climatic conditions; thus, could have different therapeutic effects. This geographical variation could mean that standardization of dose based on chemical markers is essential; however, only one study described such a standardization approach. We reported about 87% of the included studies used scientific names incorrectly. Kim and his colleagues reported similar high percentage (78.6%) in their systematic review [119]. The incorrect use of names could be the result of insufficient knowledge of taxonomy or negligence in part of researchers. The erroneous identification is a severe problem that may diminish the utility of research results. Such inappropriate uses of plant names within the literature are a permanent source of confusion for future research, search engines and databases [36].

The clinical translation of *M. charantia* L and other herbal products from the disciplines of natural product development has been slow and inefficient. The inefficiencies could be partly due to suboptimal research practices that propagate biases that hamper clinical translatability. The biases due to small studies effect, methodological flaws, use of inappropriate animal models of the human condition, use of unstandardized intervention, inappropriate use of statistics, and poor or selective reporting need immediate attention. Together, this systematic review and meta-analysis suggests that previous clinical trials of *M. charantia* L. could have been conducted based on inadequate evidence of efficacy from preclinical studies, and partly this could explains conflicting clinical trial results observed in the meta-analysis [20,120,121].

### Strength of the study

A significant strength of our study is that it is the first and timely systematic review and meta-analysis of *M. charantia* L. using animal studies. Amid a growing number of preclinical and clinical studies investigating the efficacy and underlying mechanism of action of *M. charantia* L. in glycaemic control, we provided a more in-depth insight into the current state and level of available preclinical evidence. We also provided evidence of major methodological, taxonomical flaws, and risk of bias that could potentially threat validity and clinical generalizability of preclinical studies of *M. charantia* L. Future studies could now consider improving design features that are threats to internal, constructs and external validity. Authors, reviewers, and editors should also thrive to adhere to the proposed reporting guidelines of preclinical studies such as ARRIVE [114] to improve the quality of preclinical reports and their utilization in advancing scientific knowledge.

## Conclusion

*Momordica charantia L.* reduced elevated fasting plasma glucose level with moderate quality evidence in animal models of type 2 diabetes mellitus. It also significantly reduced glycosylated hemoglobin, alanine aminotransferase, aspartate aminotransferase, alkaline phosphate, urea, serum creatinine, and several lipid profile parameters. This conclusion must be interpreted in light of strongly suspected publication bias, high risk of bias and poor methodological quality of primary studies. In order to enhance clinical generalizability, future researches should focus on standardizing dose of *M. charantia* L. with known chemical markers, provide adequate quality control data, conduct preclinical studies that are designed with random allocation, blinding of investigators and assessors, and power calculation of sample size.

## Competing interests

We wish to confirm that there are no known conflicts of interest associated with this publication and there has been no significant financial support for this work that could have influenced its outcome.

## Funding

This research did not receive any specific grant from funding agencies in the public, commercial, or not-for-profit sectors.

## Acknowledgements

This work is part of a Ph.D. thesis of ELP. World Bank supports the Ph.D. training through PHARMBIOTRAC-ACE II fellowship administered at Mbarara University of science and technology. Authors wish to thank the fellowship programme leadership for the training support.

## Authors’ contributions

Conceptualization ELP; Data curation and Formal analysis ELP, AK; Methodology ELP, AK, CDS; Project administration ELP; Supervision CDS, PBN, PEO; validation AK; Writing original draft ELP; Review the drafts for important intellectual content CDS, PBN, AK, PEO. All authors approved the final manuscript.

